# Molecular basis of calmodulin-dependent calcineurin activation

**DOI:** 10.1101/665158

**Authors:** Bin Sun, Darin Vaughan, Svetlana Tikunova, Trevor P. Creamer, Jonathan P. Davis, PM Kekenes-Huskey

**Author notes:** Corresponding authors. E-mail address (P. Kekenes-Huskey).

## Abstract

Calcineurin (CaN) is a calcium-dependent phosphatase involved in numerous signaling pathways. Its activation by Ca^2+^ is in part driven by binding of calmodulin (CaM) to a Calmodulin (CaM)-recognition motif within the phosphatase’s regulatory domain (RD); however, secondary interactions between CaM and the Calcineurin (CaN) regulatory domain may be necessary to fully activate CaN [1]. Specifically, it has been shown that the CaN regulatory domain folds upon CaM binding and that there is a region C-terminal to the canonical CaM-binding region, the ‘distal helix’, that assumes an *α* helix fold and contributes to activation [1]. We hypothesized in Dunlap *et al* [1] that this putative *α* helical distal helix is capable of binding CaM in a region distinct from the canonical CaM binding region (CaMBR) site, whereby CaN is activated. To test this hypothesis, we utilized molecular simulations including replica-exchange molecular dynamics, protein-protein docking and computational mutagenesis to model distal helix conformations. From these simulations we have isolated a potential binding site on CaM (site D) that facilitates moderate affinity inter-protein interactions that may attenuate CaN auto-inhibition. Further, molecular simulations of the distal helix A454E mutation demonstrated weakened distal helix/CaM interactions that were previously shown to impair CaN activity. K30E and G40D mutations of CaM at site D presented similar decreases in binding affinity predicted by simulations. The prediction was correlated with a phosphatase assay in which these two mutants show reduced CaN activity. This study therefore provides a potential structural basis for the role of secondary CaM/CaN interactions in mediating CaN activation.

## 1 Introduction

Calcineurin (CaN) is a phosphatase that contributes to gene expression in response to changes in Ca^2+^ homeostasis. As such, it plays integral roles in physiological processes including neurological development and maintenance, immune responses and tissue remodeling [2]. CaN is activated by rising intracellular Ca^2+^ levels. It presents modest catalytic activity in response to Ca^2+^ alone, but optimal phosphatase activity occurs upon binding Ca^2+^-saturated CaM. CaN is heterodimeric protein consisting of two domains: chain A (57-61 kDa) contains the protein’s catalytic site, while chain B (19 kDa) contributes to enzyme regulation [2, 3]. At depressed Ca^2+^ levels, the enzyme is inhibited by its auto-inhibitory domain (AID) that directly binds to the phosphatase catalytic site. Maximal relief from auto-inhibition occurs upon the binding of CaM to CaN’s regulatory domain.

Our current understanding of the protein’s activation and enzymatic activity has been shaped by a number of atomic resolution structures of CaN determined by X-ray crystallography [4–9] and nuclear magnetic resonance spectroscopy [10]. Of the many CaN structures that have been deposited to the Protein Data Bank are structures that have revealed the protein’s auto-inhibited state (PDB ID: 1aui [4]), a potentially non-physiological 2:2 CaM/CaN stoichiometric configuration [7, 11, 12], and complexes of the enzyme with immunosuppressants [5, 8] and transcription factors [6, 9]. However, much less is known about the structural basis of CaM-dependent regulation of CaN, as atomic resolution CaM/CaN complexes are limited to the intact CaM bound to small peptides comprising the CaM binding region (CaMBR) of the CaN regulatory domain [13]. From those structures, it is clear that the CaM binding region (CaMBR) assumes *α* helical secondary structure when bound to CaM. Nevertheless, the paucity of structural information inclusive of complete CaM and CaN proteins leaves critical details of CaM-dependent CaN regulation unresolved.

Other experimental studies, however, have shed light on structural mechanism of CaN activation that have eluded crystallographic and NMR probes. It is increasingly understood that CaM-dependent CaN activation depends on structural properties of the 95-residue (≈10 kDa) CaN regulatory domain [14]. This segment is intrinsically disordered [4, 13–15], which signifies that it does not assume a well-defined fold in solution and for this reason has stymied efforts to determine its structure. Nevertheless, indirect probes of its conformational properties in the absence and presence of Ca^2+^-activated CaM that have revealed important clues about the mechanism of CaN regulation. It was first observed via circular dichroism (CD) by Rumi-Masante *et al* that upon CaM’s binding, nearly fifty residues of the regulatory domain (RD)s folded into *α* helices and only half of which could be accounted for by the CaMBR region. By using hydrogen exchange mass spectrometry (HXMS), they further identified the region outside the CaMBR that gained *α*-helicity upon CaM’s binding [14]. Dunlap *et al* [1] further confirmed the observation in a mutagenesis study of that region. This region was coined the ‘distal helix’ region (DH, residue K441 - I458). Moreover, they revealed that single point mutations of three alanines within the distal helix region into glutamic acids disrupted helix formation. Importantly, these mutations reduced CaN’s apparent affinity for a substrate, p-nitrophenyl phosphate (pNPP), that competes with the AID for the CaN active site [1].

Simulations of CaN have helped bridge experimental probes of its phosphatase activity [3, 16, 17] with static, atomistic-resolution structural data. Li *et al* reported slight conformational changes of the CaN B domain following Ca^2+^ binding via molecular dynamics (MD) simulation and proposed that conformational similarity between the apo- and holo-CaN B-domain states enables the former to regulate CaN activity independent of Ca^2+^ [18]. Harish *et al* utilized virtual screening and MD simulations to design inhibitory peptides of CaN using the native AID peptide as template [19]. Simulations have also been used to study the involvement of CaN residues outside of its catalytic domain in the binding and anchoring of inhibitory immunosuppressant drugs and analogs there of [20–23]. Similarly, computational studies examining structural mechanisms of CaM-dependent regulation of targets including CaN have emerged recently, including myelin basic protein (MBP) [24] and myosin light chain kinase (MLCK) [25, 26]. In complement to these studies, we have additionally shown via molecular dynamics and Brownian dynamics simulations that the CaMBR is highly dynamic in solution in the absence of CaM, that CaM binding to the CaMBR is diffusion-limited, and that corresponding association rates are tuned by the charge density of the CaN peptide [27]. Despite these contributions, the sequence of molecular events that follow CaMBR binding and culminate in relief of CaN auto-inhibition remain unresolved.

These observations formed the basis of a working model of CaN activation whereby the folding of the intrinsically-disordered distal helix into an *α* helix-rich structure is coupled to relieving CaN autoinhibition. However, it is still not known whether the distal helix directly binds to CaM, and if so, where they might share protein-protein interaction (PPI) interfaces or how those PPIs are potentially stabilized. In large part, the challenge in identifying potential PPI sites arises because such interaction sites generally assume large, flat surfaces lacking specific interaction patterns [28], such as grooves formed between *α* helical bundles [29, 30]. Computational protein-protein docking engines have begun to address this challenge, including ZDOCK [31] and RosettaDOCK [32], which have been used to successfully elucidate structural details of intrinsically disordered peptide-involved regulation. For example, Hu *et al* utilized ZDOCK to successfully predict the binding modes between disordered Yersinia effector protein and its chaperone partner [33]. Schiffer *et al* explored the molecular mechanism of ubiquitin transfer starting from top-ranked ZDOCK predicted binding pose between ankyrin repeat and SOCS box protein 9 (ASB9) and creatine kinase (CK) [34]. Bui *et al* reported that phosphorylation of the IDP fragment of transcription factor Ets1 leads to more binding-competent structures to its coactivator as evident by MD and RosettaDOCK [35].

In this study, we utilized computational methods including protein-protein docking, enhanced sampling and classical molecular dynamics (MD) simulations to identify potential interaction sites between the distal helix and CaM. The protein-protein docking yielded several candidate interaction sites that we defined as sites A through D (Fig. 2(a)). Of these, site D on the CaM solvent-accessible surface appears to stabilize the distal helix by moderate-affinity intermolecular interactions. Among the intermolecular interactions stabilizing this putative PPI are two residues, lysine (K30) and glycine (G40) found on the ‘back-side’ of CaM distal to where CaMBR is known to bind. Their mutation to K30E and G40D were found to abolish enzyme activity [36] in another globular CaM target, Myosin Light Chain Kinase (MLCK), that apparently relies on still unresolved secondary interactions to initiate catalysis [37, 38]. Analogously, our simulations of CaM K30E and G40D variants indicate that the mutations substantially impair distal helix binding at site D. In complement to these simulations, we demonstrate that the distal helix A454E variant also destabilizes the distal helix/site D interaction in agreement with reduced phosphatase activity shown by Dunlap *et al* [1]. Our data strongly suggest that the site D and CaN distal helix region are important to CaN activation, as site directed variants at site D residues K30 and G40 reduces CaN-dependent dephosphorylation of pNPP. Based on these results, we provide an updated structural model of CaN activation by CaM that reflects specific CaM/distal helix interaction sites (see Fig. 1).

**Figure 1:**
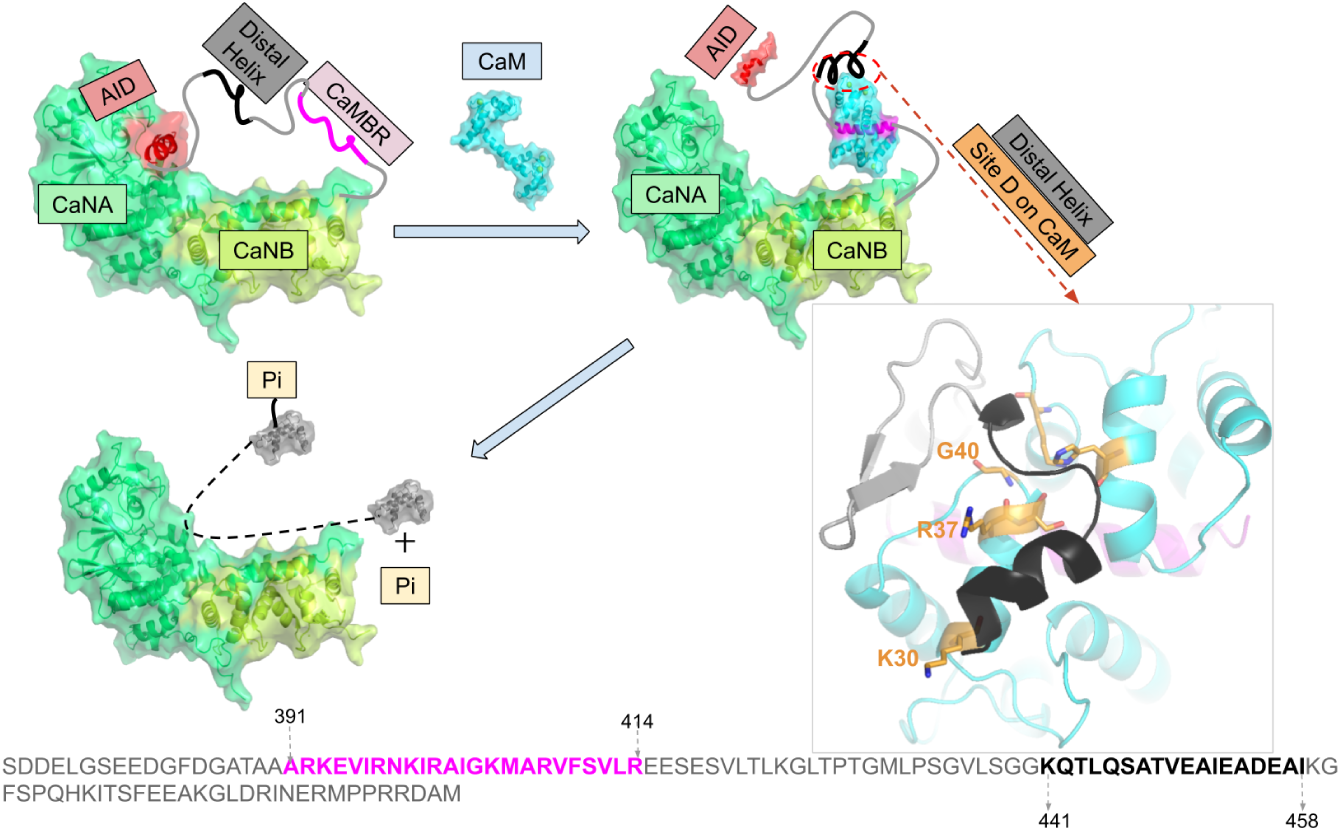
Refined model of Calcineurin (CaN) activation by Calmodulin (CaM) through direct binding of the ‘distal helix’ to CaM, based on the mechanism initially proposed in [1]. The two chains of CaN (CaNA and CaNB) are colored in limegreen and lime, respectively. AID is colored in red. CaM is colored in cyan, CaMBR is colored in magenta. The amino acid sequence of CaN RD is shown at the bottom of the panel with CaMBR and the distal helix region colored in magenta and black, respectively. In the absence of CaM, CaN is inhibited by its auto-inhibitory domain (AID). After CaM binds the CaM binding region (CaMBR) on the CaN regulatory domain, a secondary interaction between CaM and a ‘distal helix’ ultimately remove the AID from the CaN catalytic domain. The activated CaN enzyme catalyzes the dephosphorylation of target proteins essential to myriad physiological functions.

## 2 Methods

Our simulation protocol consisted of four primary steps. 1) replica exchange molecular dynamics (REMD) simulations of the isolated CaN distal helix region were used to generate trial conformations for protein/protein docking with CaM. 2) The ZDOCK protein/protein docking engine yielded initial poses for putative CaM/CaN interaction sites. 3) The docking poses were refined using extensive, microsecond-length molecular dynamics simulations. 4) Binding affinities based on MM/GBSA were used to rank-order distal helix/CaM poses for sites A-D. We further challenged the predicted structural models by introducing mutations in the distal helix and putative interaction site D that have been experimental probed in prior works. We describe these steps in detail below and additionally summarize pNPP phosphatase assay.

### 2.1 Replica exchange molecular dynamics (REMD) sampling of the isolated distal helix

In accordance with our approach in [27], we performed replica exchange molecular dynamics (REMD) simulations of the distal helix region (K441-I458) in the absence of CaM to exhaustively sample likely conformational that are in equilibrium. The distal helix peptide was constructed by the auxiliary tleap program in Amber16 [39] in an extended configuration and parameterized using the Amber ff99SBildn [40] force field. The peptide was then minimized via sander [41] *in vacuo* until convergence of the energy gradient (drms ≤ 0.05) or the number of steps 1 × 10^5^ (with first 50 steps of steepest decent and rest steps of conjugate gradients algorithm) was satisfied. The minimized structure was then used as the starting structure for REMD simulations coupled with the Hawkins, Cramer, Truhlar pairwise generalized born implicit solvent model [42] via the *igb = 1* option in Amber. The monovalent 1:1 salt concentration was set to 0.15 M and a non-bound cutoff of 99 Å was chosen. Ten replicas were created with temperature ranges spanning 270-453 K. The temperature of each replica was calculated via the Patriksson *et al* webserver [43] to ensure the exchange probability between neighbouring replicas was approximately 0.4, as recommended in [44, 45]. Each replica was first subjected to 1 × 10^5^ steps of energy minimization via pmemd with the first 50 steps being steepest decent and rest steps being conjugate gradients. The minimized systems were subsequently heated from 0 to their respective target temperatures over an 800 ps interval using a timestep of 2 fs with Langevin thermostat. The equilibrated replicas were then subjected to 100 ns of production REMD simulations under target temperature with Langevin thermostat. The SHAKE [46] algorithms were used for REMD simulations. Clustering analysis with hierarchical agglomerative (bottom-up) approach using cpptraj were conducted on the 300 K REMD trajectory to divide the trajectory into ten clusters; the average root mean squared deviations (RMSD) between each cluster was around 6 Å.

### 2.2 Docking of distal helix to CaM/CaMBR complex via ZDOCK

The protein-protein docking webserver ZDOCK3.0.2 [31] was used to determine probable binding poses for the REMD-generated distal helix conformations on the CaMBR-bound CaM complex. The CaM/CaMBR complex configuration was obtained from the Protein Databank (PDB ID: 4q5u [47]). It has been reported that 62% percent of experimentally-resolved PPIs are characterized by the binding of an *α*-helical peptide within grooves formed between adjacent *α*-helical on the target protein surface [30]; therefore we narrowed the ZDOCK search to four *α*-helical-containing regions on the CaM solvent-exposed surface. These sites are shown in Fig. 2(a), from which we determined a list of probable amino acid contacts as input to ZDOCK (see Table S1). During the ZDOCK calculations, the receptor (CaM/CaMBR complex) was kept fixed while grids were constructed around receptor with the size being 80*X*80*X*80 and spacing being 1.2 Å. The ligand (distal helix) was then docked via fast fourier transform (FFT) algorithm on the 3D grids. The scoring function consists of interface atomic contact energies (IFACE) [48], shape complementarity and electrostatics with charge adopted from CHARMM19 force field [49]. The initially generated 2 × 10^3^ poses were subjected to a culling process to eliminate those having no contacts with residues we specified in Table S1. After culling, there were zero, two, eighty-eight and three poses left at sites A-D, respectively. For case in site A, ZDOCK then instead output all initial 2 × 10^3^ poses and we found that the top ten poses were still near site A (Fig. 2(b)). The pose with highest score at each site was chosen for further refinement using molecular dynamics.

**Figure 2:**
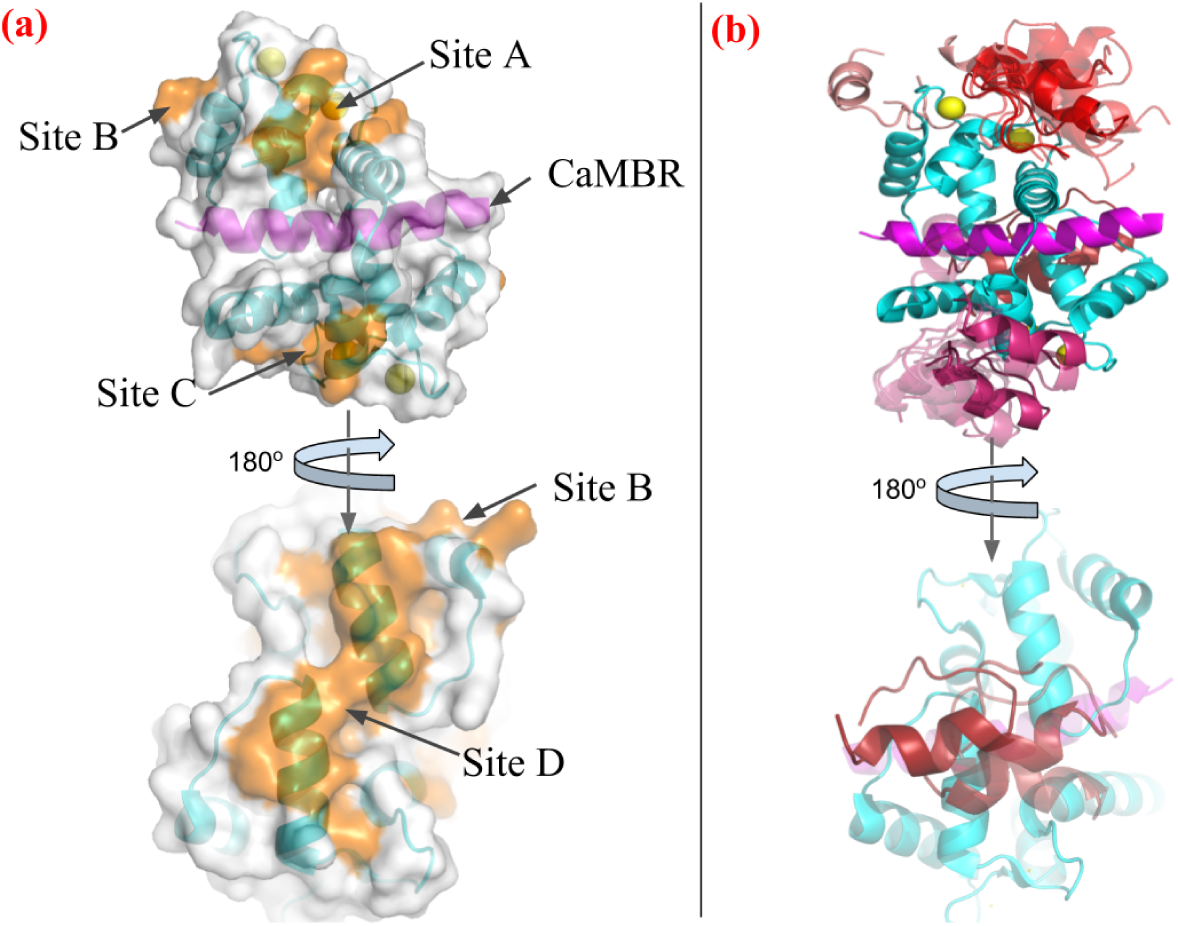
(a) Four tentative binding sites (orange) on the surface of CaM-CaMBR complex. CaM is colored in cyan, CaMBR is colored in magenta and Ca^2+^ ions are colored in yellow. (b) ZDOCK predicted conformations of distal helix interacting with CaM/CaMBR complex at each site. Predicted distal helix conformations from site A to D are colored as red, salmon, warmpink and firebrick, respectively. /net/share/bsu233/CaMDH/zdock_MD/esitmate_SASA/log.pml

### 2.3 Conventional molecular dynamics (MD) simulations of ZDOCK-generated distal helix/CaM poses

Explicit-solvent MD simulations were performed on the ZDOCK-predicted distal helix/CaM complexes to further refine the distal helix binding poses. The amino acid sequence from CaMBR to distal helix is shown at the bottom of Fig. 1 and the sequence definition of CaMBR and distal helix are the same as [1]. We first inserted peptide linkers for each pose between the CaMBR C-terminus (R414) and the N-terminus (K441) of the distal helix via tleap. The initial linker was generated via tleap and energy-minimized as done in Sect. 2.1. The minimized structures were subsequently simulated *in vacuo* to heat the systems from 0 to 300 K. The last frame of the short equilibration run was subject to additional energy minimization *in vacuo* to facilitate its compliance with the distal helix and CaMBR termini. The top poses from ZDOCK presented distal helix orientations that were all compatible with the CaMBR and linker configurations. The optimized linker was placed adjacent to the CaMBR and distal helix; tleap was then used to link the peptide components. The resulting structures were then subjected to energy minimization, followed by a 100 ps heating process to raise the system temperature to 300 K for which all atoms except the linker were fixed via the ibelly function in sander MD engine of Amber. This minimization and heating was perform *in vacuo* and the purpose is to further relax the linker in the presence of distal helix and CaM/CaMBR complex. The last frame of heating stage was used as input configurations for explicit-solvent molecular dynamics simulations.

Each *in vacuo* starting configuration was solvated in a TIP3P [50] waterbox with 12 Å boundary margin. K^+^ and Cl*^−^* ions were added to neutralize the protein and establish a 0.15 M salt concentrations. After parameterizing the system using the ff14SB force field [51] via tleap, the system was subjected to an energy minimization, for which all atoms except hydrogens, water and KCl ions were constrained by the ibelly functionality. The cutoff value for non-bond interactions was set to 10 Å. A 2 fs timestep was chosen, as SHAKE [46] constraints were applied on bonds involving hydrogen atoms. Two heating procedures were performed to heat the system from 0 to 300 K using the Amber16 sander.mpi engine [41]. In the first heating stage, the *ibelly* function was used to keep all system except the water box and KCl ions fixed. The water box was heated to 300 K over a 100 ps interval under the NVT ensemble. For the second heating stage, the entire system was heated from 0 to 300 K over 500 ps under the NPT ensemble, for which the backbone atoms of CaM, CaMBR and distal helix were constrained by an harmonic potential (force constants of 3 kcal mol*^−^*^1^ Å*^−^*^2^). Thereafter, an additional 1 ns equilibrium stage was conducted at 300 K under the same constraints, but with a reduced force constant of 1 kcal mol*^−^*^1^ Å*^−^*^2^. These equilibrium simulations were followed by 100 ns production-level MD simulations. The weak-coupling thermostat [52] was used during the simulation. Clustering analysis was performed on the production trajectory using the same strategy in Sect. 2.1. The average root mean squared deviations (RMSD) between each cluster was around 6 Å. Based on the rationale that extending simulations using less-frequently sampled structures provides greater overall sampling of the conformational space [53], we identified 5-6 low-probability states as inputs for subsequent MD simulations. Approximately 1 *µ*s of trajectory data were simulated in total for each site.

### 2.4 MD simulations of CaM (K30E and G40D) and CaN distal helix variants (A454E)

Clustering analyses is a common used data-mining method in MD simulation which groups trajectories into clusters based on structural similarity [54]. In this study, clustering analyses was performed on the production-level MD trajectories of the distal helix/CaM configurations that yielded the most favorable binding scores by MM/GBSA. A representative structure of the most populated cluster was selected as an input for *in silico* mutagenesis in order to validate the model against experiment. Namely, the CaM K30E and G40D variants, as well as the CaN A454E variant, were built by replacing and regenerating the amino acid side chains using tleap. The resulting structures were energy minimized with a stop criterion of (drms <= 0.05) for the energy, during which all atoms except the mutated residues were fixed via the ibelly function in Amber. The energy-minimized structure was then solvated and simulated according to the same procedure in Sect. 2.3. All simulation cases in this study were listed in Table S3.

The binding free energy between distal helix and CaM was estimated via Molecular Mechanics-Generalized Born and Surface Area continuum solvation (MM-GBSA) [55].

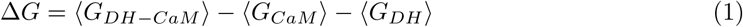

Where 〈*G_DH−CaM_*〉, 〈*G_CaM_*〉 and 〈*G_DH_*〉 are ensemble-averaged free energies of distal helix-CaM complex, CaM and distal helix, respectively. In this study, the trajectories of these three components were extracted from MD trajectories via cpptraj at a 2 ns frequency. The generated sub-trajectories were used as input of MMPBSA.py script shipped with Amber16 to calculate the free energies of each part. The salt concentration was set as 0.15 M with generalized Born model option setting as *igb = 5*. No quasi-harmonic entropy approximation was made during the calculation.

### 2.5 Structural Analyses

Clustering analysis, root mean squared deviations (RMSD)/root mean squared fluctuations (RMSF) calculations, hydrogen bonds and secondary structure analysis were performed via cpptraj [56] shipped with Amber16. The reference structure of RMSD calculation was CaM/CaMBR crystal structure (PDB ID: 4q5u [47]). Prior to RMSF calculation, the whole MD trajectory was first rms fitted to the first frame using CaM/CaMBR backbone atoms. Secondary structure probability of each residue was calculated using cpptraj with define secondary structure of proteins (DSSP) algorithm [57]. The colvar module [58] within VMD was used to assess the total *α*-helix content of REMD-generated distal helix and DH_A454E_ conformation. The *hbond* command within cpptraj was used to analyze hydrogen bonds between distal helix and CaM/CaMBR. During the hbond analysis, the angle cutoff for hydrogen bonds was disabled while the default 3 Å cutoff between acceptor and donor heavy atoms was used. Scripts and cpptraj input files used for above analyses will be publicly available at https://bitbucket.org/pkhlab/pkh-lab-analyses/src/default/2018-CaMDH.

### 2.6 Calcineurin phosphatase assay using para-nitrophenyl phosphate (pNPP) substrate

#### Materials

pNPP was obtained as the bis(tris) salt (Sigma), dithiothreitol reducing agent (Sigma), assay buffer (80mM Tris pH 8, 200mM KCl, 2mM CaCl_2_), MnCl_2_. *Preparation of Enzymes and Proteins*. The wild-type, K30E, and G40D variants were obtained from JP Davis and prepared at a concentration range of 300 *µ*M to 3 mM in assay buffer. *Enzyme Assay*. Phosphatase assays were performed using 30 nM calcineurin, and 90 nM calmodulin in 96-well Corning Costar microtiter plates with a reaction volume of 120 *µ*L. Assays proceeded in the manner of [1] with each CaM assayed in triplicate and over three plates to account for technical variation. Control reactions absent calcineurin were added to the end of each lane with 200 mM pNPP to determine the rate of enzyme-independent substrate hydrolysis. *Kinetic Analysis*. The pNPP substrate reactions were varied over 11 concentrations, increasing from 0 mM to 200 mM for each column. 60 minute UV-Vis recordings were obtained on a Molecular Devices FlexStation 3 plate reader using Softmax Pro 7 software at 405 nm with 10 minute read intervals. The resulting data was inspected for appropriate Michaelis-Menten kinetics, such as proper substrate saturation, by plotting absorbance against substrate concentration. Following this, the readings were linearized to produce the double reciprocal Lineweaver-Burk plot for extraction of *V_max_* and *K_M_* based on the following equation:

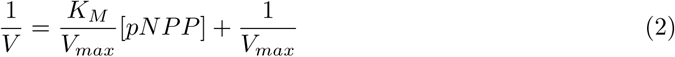

## 3 Results

Prior studies [14, 59] have indicated that CaM binding to the CaN’s canonical CaM-binding region requires secondary interactions beyond this region to fully activate the phosphatase. Rather, CaN activity is likely dependent on a secondary interaction between the CaN regulatory domain and CaM. A study by Dunlap *et al* [1] suggested that a distal helix region spanning residues K441 to I458 was likely involved in CaM binding. However, it was unclear which region(s) of the CaM solvent-exposed surface would contribute to a potential PPI. We therefore used molecular dynamics and protein-protein docking simulations to identify plausible wild-type CaN interaction sites on CaM, and challenge these predictions with mutagenesis. Our predicted site was validated using a CaN pNPP phosphatase assay.

### 3.1 Regulatory domain (RD)-construct propensity for secondary structure formation in absence of CaM

Circular dichroism (CD) and HXMS analysis in [14] suggest that there exists *α*-helical structure beyond the canonical CaMBR region after CaM’s binding. We therefore sought to assess *α* helicity in the REMD-simulated distal helix peptides. Previously [27], we found that extensive MD simulations of the isolated CaMBR yielded a small population of *α*-helical structures suitable for binding CaM in its canonical binding pose [60]. We therefore applied a similar REMD procedure (see Sect. 2.1) to the proposed distal helix segment of the CaN regulatory domain to assess the propensity for the spontaneous formation of secondary structure in the absence of CaM. Here, we performed 100 ns of REMD simulations on the wild-type (WT) distal helix as well as a A454E variant. The latter was considered as it has been reported to exhibit reduced *α*-helical content in the presence of CaM [1], which is suggestive of abolishing the distal helix/CaM interaction. Following the REMD simulations, we performed clustering analysis to identify the predominant conformations of the two peptide configurations. Interestingly, we observed that both the WT distal helix and its A454E mutant partially fold into an *α* helix in the absence of CaM. As shown in Fig. 3(a), the representative structure of most populated clusters of the distal helix and A454E mutant (83.8% and 85.3% of the total trajectory, respectively) both contain helical fragments. While the overall *α*-helix contents (≈ 45%) of these two fragments were statistically indistinguishable, a contiguous helix was formed in the WT distal helix, whereas it was fragmented in the mutant. These helicity features are further quantified as residue’s *α*-helix structural probability shown in Fig. 3(b): the distal helix region has the maximum probability present at middle region while the A454E has maximums present near the two terminis. Both the simulated distal helix and its variant therefore could adopt *α*-helix content in the absence of CaM, but it remains to be determined whether the dominant structures are capable of binding the CaM surface. We note that experimental assays of the complete RD do not detect significant secondary structure; this discrepancy may be a result of using substantially different RD lengths (S374 to Q522 residues in Rumi-Masante *et al* [14] and K441-I458 in this study). We discuss this difference in further detail in Limitations (Sect. 4.5).

**Figure 3:**
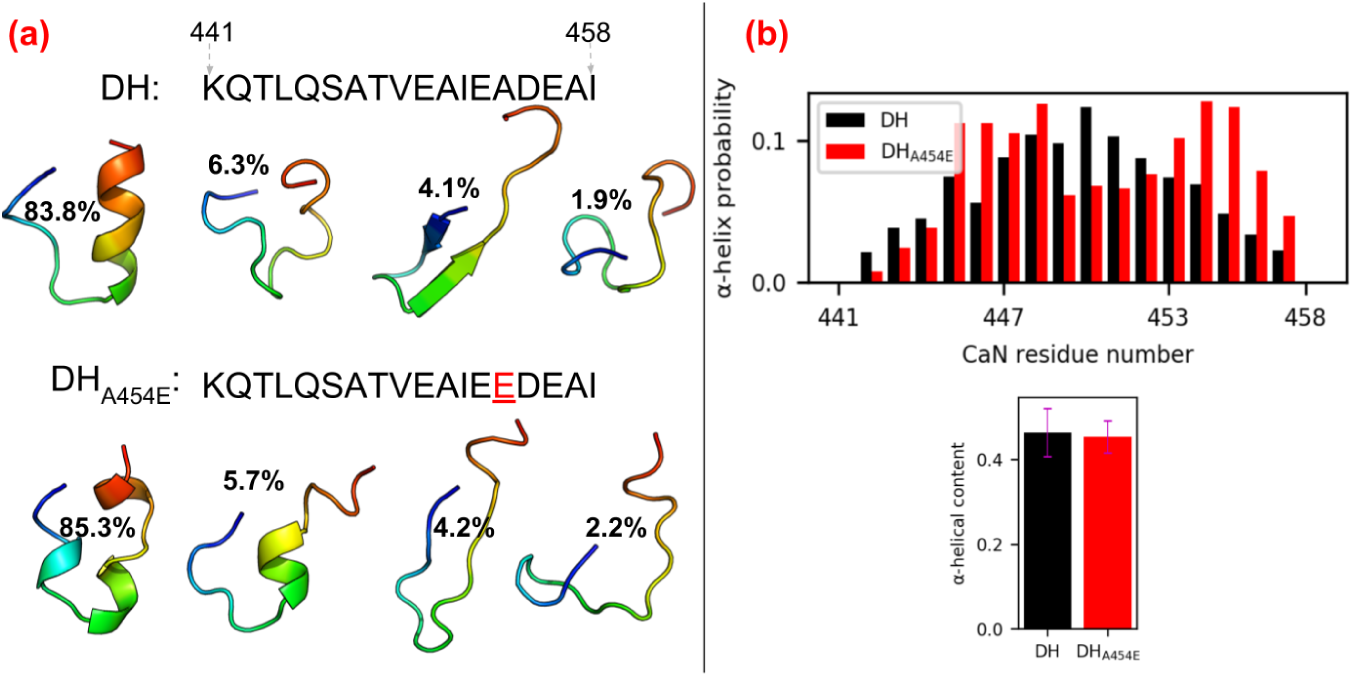
(a) Sequence of distal helix/DH_A454E_ and representative structures of four most populated clusters from 100 ns REMD simulations. The structures are colored in rainbow with N-termini as blue and C-termini as red. (b) Secondary structure probability of each residue calculated from REMD trajectory via cpptraj with DSSP algorithm. The lower panel shows the total *α*-helix contents of two fragments calculated via the colvar module of VMD. https://bitbucket.org/pkhlab/pkh-lab-analyses/src/default/2018-CaMDH/rmsf.ipynb

### 3.2 Protein-protein interactions between RD-construct and peptide-bound CaM

The overwhelming majority of CaM-containing complex structures resolved to date include only limited fragments of the bound target protein [60]. CaM-bound CaN is no exception, as the mostly likely physiological conformation [47] consists of monomeric CaM in a canonical ‘wrapped’ conformation about a target region in CaN(A391-R414) [13]; however, it is evident that secondary interactions beyond this domain play a role in CaN activity, yet atomistic-level structural details of these interactions have not yet been resolved. Therefore, in order to resolve potential binding regions for the distal helix region, we seeded a protein-protein docking engine, ZDOCK [31], with candidate *α*-helical structures identified through REMD simulations. The docking simulations were performed in regions that included grooves formed between *α* helices we identified at the CaM solvent-accessible surface. We selected these regions, since such secondary structures are believed to nucleate protein-protein interactions [61]. Furthermore, a thorough examination of protein-protein complex structures in the Protein Data Bank in 2011 suggested that *α* helices contribute to 62% of all PPI interaction surfaces [30] between binding partners. Narrowing the search region on CaM to those containing *α*-helical regions yielded four candidate sites (A-D) that spanned nearly the entire CaM solvent-exposed surface (see Fig. 2(a)).

The most energetically-favorable distal helix/CaM poses predicted via ZDOCK at sites A-D are summarized in Fig. S1. The docked poses reflect significant interactions of at least the distal helix C-terminal loop with loops bridging adjacent *α*-helices on the CaM surface. At site A, polar residues near N97, Y99 and D133 from two of the C-terminal CaM domain’s loops interact with the distal helix, compared with just one EF-hand motif loop at site B (D129, D133 and D135). The site C poses were primarily stabilized by hydrophobic interactions formed from CaN residues L444/I458 and F16/L4 on CaM, in addition to a loop-loop interaction via CaM D64. The site D poses reflected distal helix C-terminal loop interactions with CaM EF-hand loop residues near N42 and K94. Most poses were unsurprisingly parallel to *α*-helical/*α*-helical ‘grooves’ on the CaM solvent-exposed surface and were evidently anchored through interactions between the proteins’ loop regions.

In contrast, we found that the A454E variant docked poorly at sites A-D (see Fig. S2), as assessed by the proximity of docked poses to the designated CaM sites in fact, most predicted poses tended to localize toward site A, albeit with weak interactions. Moreover, we speculate that the impaired binding of DH_A454E_ may arise from its fragmented *α* helical structure, in contrast to the contiguous regions for the WT variant (see Table S2 for docking scores and Fig. S1/Fig. S2 for docking poses). Although docking scores were provided by the ZDOCK algorithm to rank order potential poses, we did not analyze these scores in detail as we later refined these structures using more detailed simulations and energy expressions. This refinement corrects for artifacts from the ZDOCK algorithm, which assumes rigid conformations for both proteins that would ordinarily be expected to relax in the bound complex. Hence, in the following section we pursue extensive microsecond-scale all-atom MD simulations to refine and assess the predicted poses.

The docked CaN/CaM configurations from the previous section were intended as inputs for MD-based refinement of nearly intact CaN regulatory domain complexes with CaM. Subsequent refinement using microsecond-length MD simulations relax the rigid protein conformations assumed in ZDOCK. To refine these poses, we linked the docked distal helix fragments with the CaMBR fragment resolved in the CaM/CaN complex (PDB ID: 4q5u) from [47]. Each of the four candidate binding sites yielded distal helix orientations that were compatible with the 26 residue-length linker. Following initial optimizations of the linker described in Sect. 2.3, we performed *µ*s-length, explicit solvent simulations with the regulatory domain bound to CaM. Since the predicted A454E distal helix poses appeared to be inferior to those of the WT variant, we refined only the WT poses and thereafter introduced A454E mutations to the refined conformations.

We first assess the integrity of the predicted binding modes based on Molecular Mechanics-Generalized Born and Surface Area continuum solvation (MM-GBSA). MM/GBSA scoring of the MD-generated configurations provides a coarse estimate of binding affinity without significantly more expensive free energy methods. We reported the binding free energy of distal helix between CaM/CaMBR as well as between CaMBR and CaM in Fig. 4. Significantly, we found that binding of WT distal helix at the CaM site D yielded a more pronounced favorable average binding free energy (Δ*G* ≈ −27.7 kcal mol*^−^*^1^) than sites A, B and C (−2.5 kcal mol*^−^*^1^, −17 kcal mol*^−^*^1^, −22.5 kcal mol*^−^*^1^) with P-values (1 × 10*^−^*^4^, 2.8 × 10*^−^*^3^ and 1.144 × 10*^−^*^1^, respectively) confirming that the means are significant compared to the null hypothesis. Similarly, the binding free energies of distal helix interactions were generally substantially weaker (−2.5 to −27.5 kcal mol*^−^*^1^) than those between the CaMBR and CaM (Δ*G <* −1.20 × 10^2^ kcal mol*^−^*^1^) Although MM-GBSA is a very approximate scoring method for molecular complexes, the consistent trends in numbers of hydrogen bond contacts, RMSF amplitudes and binding scores suggests that the site D is the most likely region for forming stable CaM/distal helix interactions.

**Figure 4:**
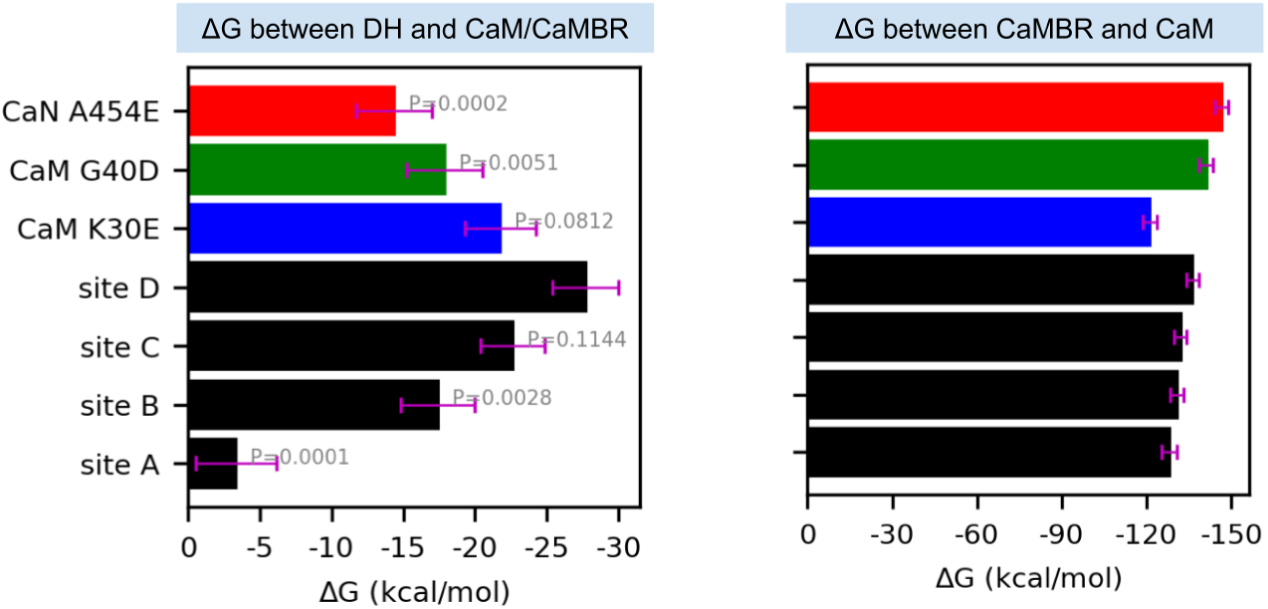
Approximate binding free energies between CaM and the distal helix (left) or CaMBR regions (right) via Molecular Mechanics-Generalized Born and Surface Area continuum solvation (MM-GBSA). Black bars correspond to wild-type CaN, whereas colored bars utilize the A454E CaN and CaM variants. The calculation was conducted on frames extracted every 2 ns from MD trajectories. The error bar represents standard error of mean. The values above bars in the left panel are P values of each case with null hypothesis that their mean values are equal to site D. https://bitbucket.org/pkhlab/pkh-lab-analyses/src/default/2018-CaMDH/mmgbsa.ipynb

We supplement the energy scores with structural indicators of stability, namely contacts and RMSF. We report in Fig. 5 the corresponding root mean squared deviations (RMSD) and root mean squared fluctuations (RMSF) of the peptide backbone atoms from CaM and CaMBR. We additionally include two CaM variants with mutations at site D, which we rationalize later in Sect. 3.3. We found that the average RMSD values of the MD-predicted conformations relative to the experimentally-determined CaM/CaMBR structure were at or below 2 Å; we attribute these small fluctuations to stable CaM/CaMBR interactions that were insensitive to the distal helix docking pose. Similar to the RMSD data, the CaM and CaMBR RMSF values are comparable in amplitude and nearly indistinguishable between distal helix/CaN docking poses, with most residues presenting values below 1.5 Å. The prominent peaks in excess of 5.0 Å correspond to the CaM termini and the N-terminus of the CaMBR. We additionally observe a variable region midway along the CaM sequence, which corresponds to the labile linker between its globular N- and C-domains that is implicated in allosteric signaling [62].

**Figure 5:**
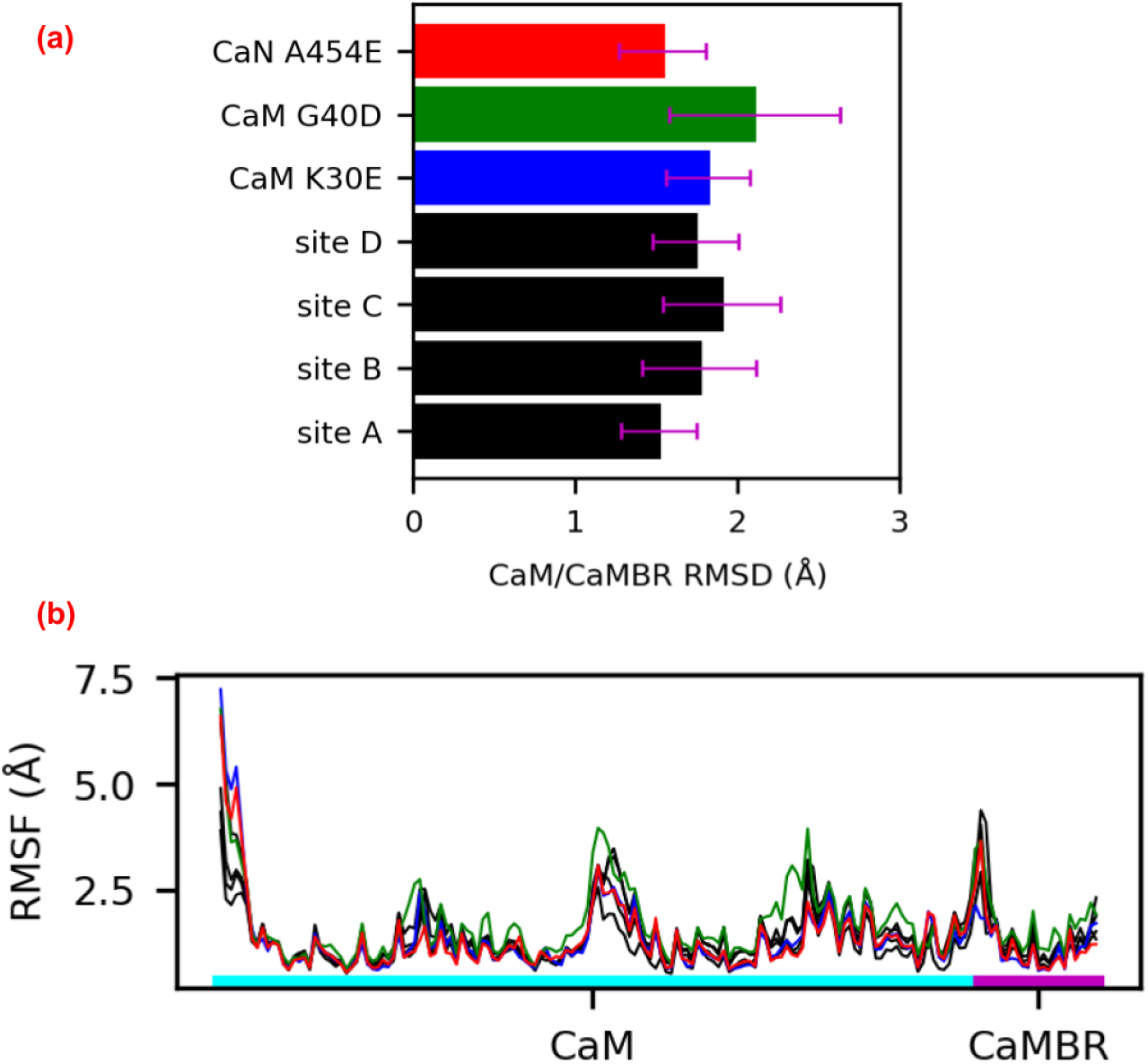
(a) Root mean squared deviations (RMSD) of peptide backbone atoms of CaM and CaMBR from *µ*s-length MD simulations. The reference structure for the RMSD calculation was based on the CaM/CaMBR crystal structure (PDB ID: 4q5u). (b) Root mean squared fluctuations (RMSF) of non-hydrogen atoms in CaM and CaMBR. https://bitbucket.org/pkhlab/pkh-lab-analyses/src/default/2018-CaMDH/rmsf.ipynb

The small and statistically indistinguishable RMSF values for the CaM/CaMBR in Fig. 5 suggest that distal helix binding had negligible impact on binding the CaM recognition motif. This is an important observation, as viable binding poses for the distal helix are expected to preserve the binding between the CaMBR and CaM. We base this assumption on CD data collected in [63] that indicated substantial alpha helical character in the CaM/CaN complex following dissociation of the distal helix domain. Therefore, we then assessed the integrity of the distal helix poses using RMSF analyses and measurements of inter-protein contacts. In Fig. 7 we report representative configurations of the distal helix region (red) in complex with CaM (cyan), as well as their corresponding per-residue RMSF values in Fig. 6. To guide interpretation, we hypothesized that RMSF values above 5 Å were indicative of poorly stabilized residues. We later rationalize this value by comparing approximate binding energies as computed by MM-GBSA. At site A, both the distal helix/CaMBR linker and the distal helix reflect RMSF values in excess of ~10 and ~15 Å, respectively. These large fluctuations arise from the breadth of binding orientations evident in Fig. 7(a), which we interpreted as a poorly-stabilized configuration. Similarly, the site B configurations also appear to be loosely bound, based on linker and distal helix RMSF values beyond 10 Å. In contrast, the distal helix RMSF values at sites C and D were below 5 Å, with the latter site reporting the smallest values among the sites we considered, which is evidence of a stable binding configuration.

**Figure 6:**
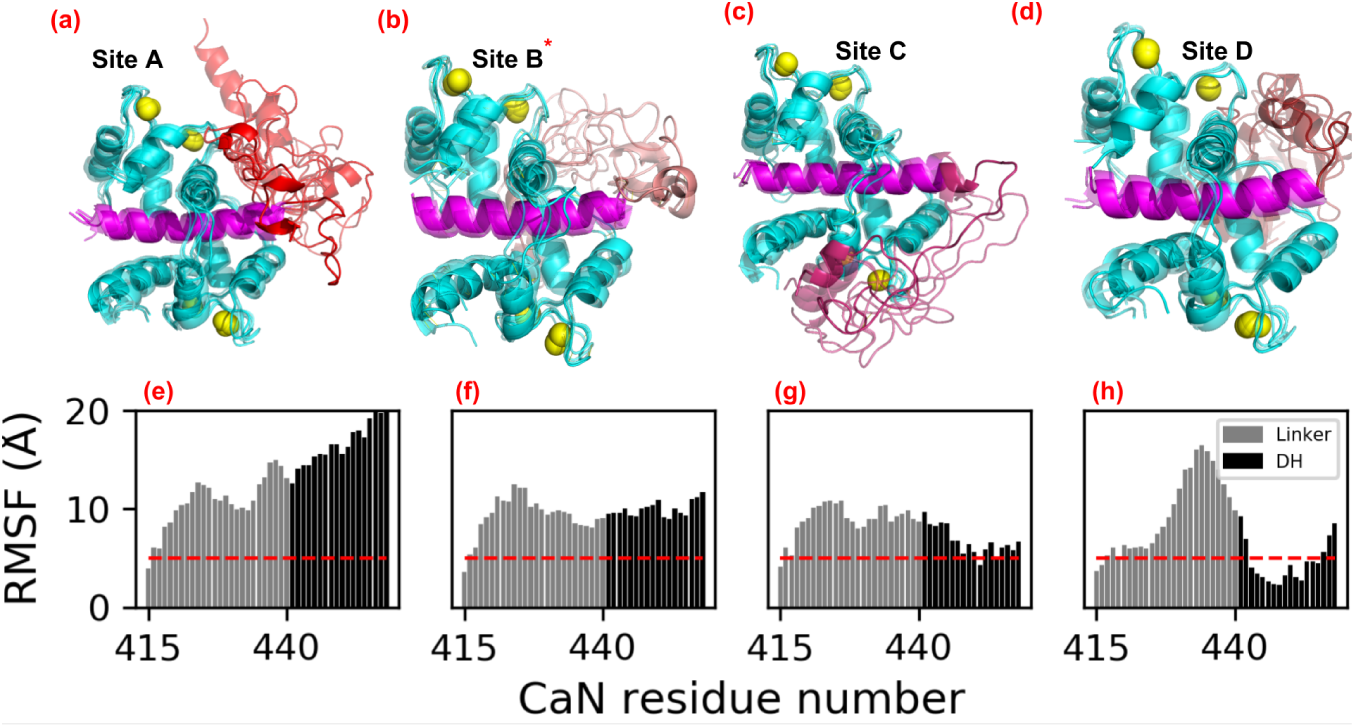
(a-d) Representative structures of from the microsecond length MD simulations initialized from ZDOCK-predicted distal helix poses. CaM is colored in cyan, CaMBR is colored in magenta and Ca^2+^ ions are depicted as yellow spheres. The linker and distal helix regions in site A-D are colored as red, salmon, warmpink and firebrick, respectively. (e-h) Non-hydrogen atom RMSFs of linker and distal helix residue calculated from MD simulations of each site, as an indicator of binding stability. The red dash line depicts RMSF value as 5 Å. * During the MD simulations, distal helix structures initiated at site B migrated toward site D (see Fig. S3). https://bitbucket.org/pkhlab/pkh-lab-analyses/src/default/2018-CaMDH/rmsf.ipynb

**Figure 7:**
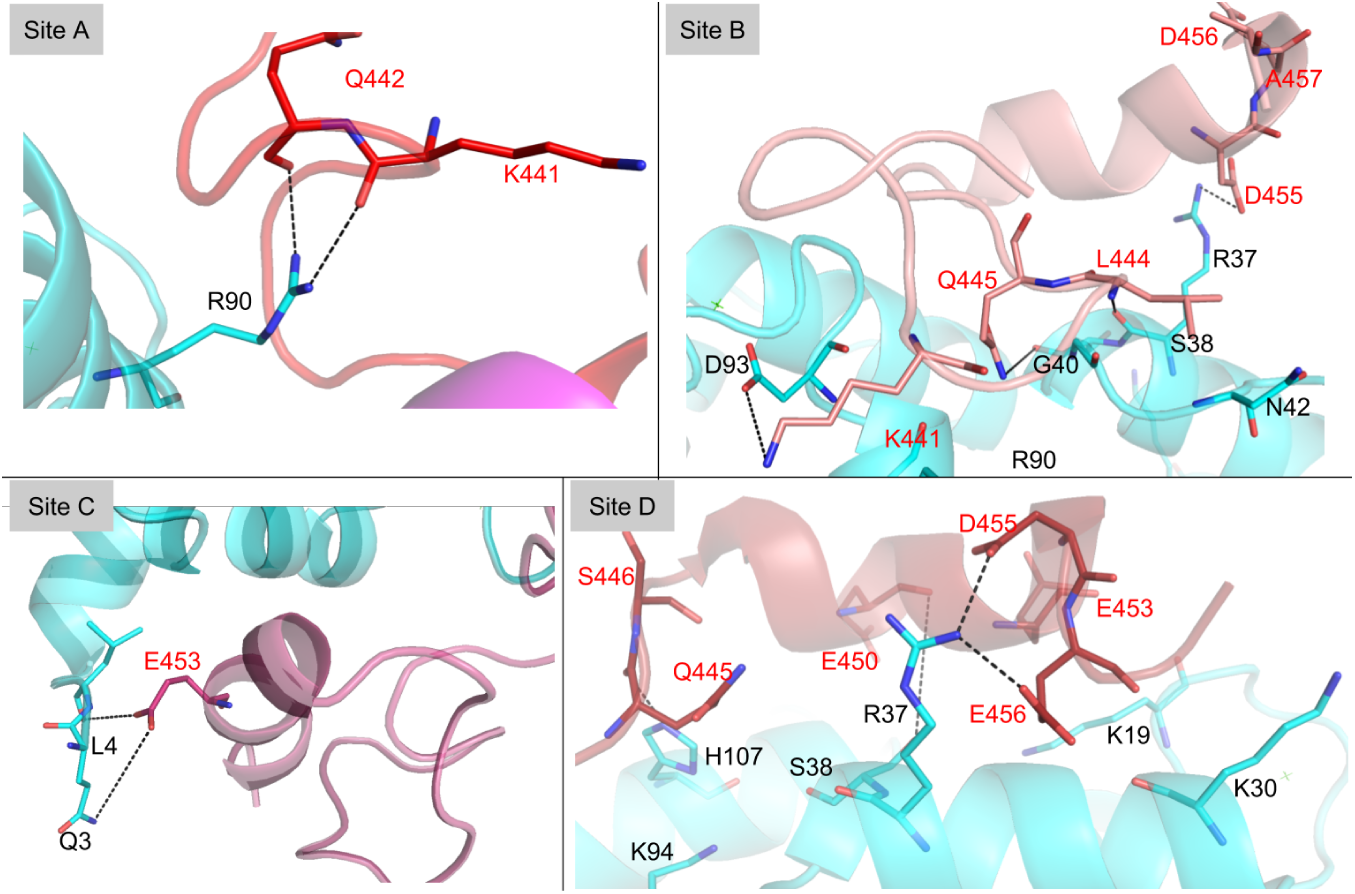
Interaction between linker/distal helix of CaN and CaM at site A-D. Key residues at the interaction surface are shown in sticks with black labels for CaM residues and red labels for distal helix residues. See Table S1.3 for specific values.

As has been shown in other proteins regulated by disordered protein domains [64–66], there are often multiple poses the contribute to regulation. We therefore assessed the most significant inter-protein contacts contributing to the ensemble of distal helix binding poses at sites A-D. Among these poses, the distal helix configurations at site D presented the lowest distal helix RMSF values among the considered sites. Significantly, the site D distal helix configuration presented several hydrogen bond-facilitated interactions with CaM, including two long-duration (37% and 55% of sampled configurations) interactions between Q445 and CaM residues R37/K94, pairing of CaM K21 with glutamic acids E453 and E450, as well as E456 with CaM residues K30 and R37. Contacts between CaM and CaN, as well as their longevities (as assessed by the percentage of MD frames satisfying a hydrogen bond contact cutoff of 3 Å between oxygen and nitrogen atoms) are additionally quantified in Fig. 8 (specific values are listed in Table S1.3). The latter data indicate a modestly greater degree of hydrogen bonding of the distal helix at site D (10 h-bonds were above 10%) versus site B (9), and a significantly greater degree relative to sites A (1) and C (3). Furthermore, the site D pose appears to be stabilized by both the N- and C-domains of CaM (residues D20-S38 and R90-N111, respectively). We speculate that this bi-dentate interaction could improve CaMBR binding by locking CaM into its collapsed configuration and thereby prevent disassembly. Although during the simulations, the distal helix at site D maintained significant *α*-helix (see Fig. S3 and Fig. S4), we note that a significant percentage of the predicted structures exhibited beta sheet character in the linker region (see Fig. S5) that was not observed in the CD cpectra collected by Rumi-Mansante *et al* [14]. This persistent secondary structure was limited to a few residues (see Fig. S5 and Fig. S6) thus may be beyond the limits of detection in earlier CD experiments. We comment on this further in the Limitations (see Sect. 4.5). Meanwhile, site B reflected interactions with both CaM terminal domains that were attenuated, while sites A and C were mostly bound by interactions of their linker regions with the CaM N-domain. Interestingly, we observed that the distal helix poses originating at site B migrated toward site D (see Fig. S3), which likely explains the higher hydrogen bonding in site B versus sites A and C.

**Figure 8:**
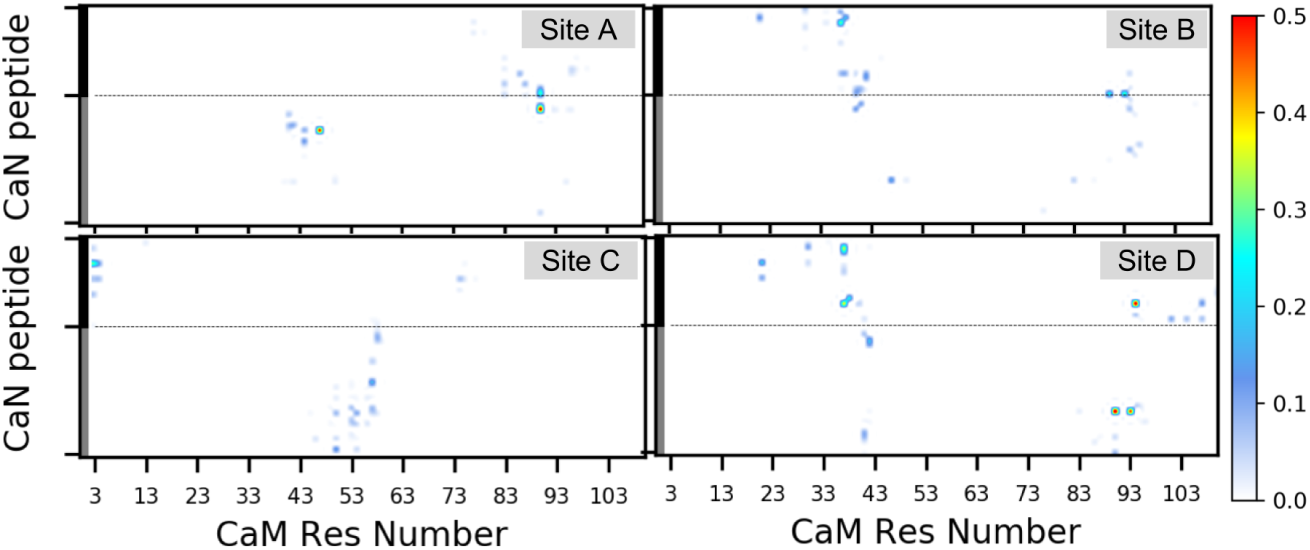
Percentage of simulated frames which have hydrogen bonds formed between CaN peptide (linker and distal helix) and CaM. The linker and distal helix are indicated by grey and black bar, respectively.

As a result of HXMS conducted by Rumi-Masante *et al* [14] of the RD construct CaN in solution with CaM, it is apparent that residues R414 through E456 are within a stretch of residues that are somewhat protected from solvent, which suggest that relief of CaN autoinhibition entails binding at least the distal helix region. We note that the HXMS data could not precisely distinguish which residues were protected, as proteolysis and mass spec was conducted on short peptides. Further, HXMS data detects only bonds involving backbone amide protons, thus we speculate that the CaN site chain interactions with CaM may stabilize the distal helix alpha helical structure. Hence, we suggest that CaM/CaN configurations that stabilize the distal helix region likely contribute to CaN activation. Based on this rationale, the small RMSF values and extensive hydrogen bonding of the CaN distal helix with the CaM site D relative to other ZDOCK identified regions suggest that CaN is most stabilized at site D.

### 3.3 Effects of putative CaN/CaM site D mutagenesis

MD simulations of the WT CaN CaMBR-distal helix sequence suggest that CaM site D is a probable binding region for the CaN regulatory domain. To challenge this hypothesis, we performed MD simulations of CaN distal helix and CaM site D variants that could reduce CaN activity to test whether the distal helix/CaM interaction was impaired. Namely, we introduced the CaN A454E and CaM K30E and G40D mutations into the MD-optimized WT structures. We elected to mutate the WT CaMBR/distal helix complexes with CaM, as the WT complex appeared to have favorable stability, whereas repeating the REMD/zdock steps with the mutants may not have yielded viable configurations. The proposed A454E CaN variant was based on CD data collected by Dunlap *et al* [1] that demonstrated reduced *α*-helical content upon binding CaM relative to the WT with impaired CaN activation. The CaM variants we examined in this study were based on experimental mutagenesis studies [36] of CaM-dependent Myosin Light Chain Kinase (MLCK) activation, for which secondary interactions beyond the canonical CaM binding motif were required for enzyme activation [37, 38] (Fig. 9(a)). Although these secondary CaM interactions are involved in directly binding the MLCK catalytic domain in contrast to CaN [37], two residues (K30 and G40) implicated in binding [36] reside within the site D identified in our simulations.

**Figure 9:**
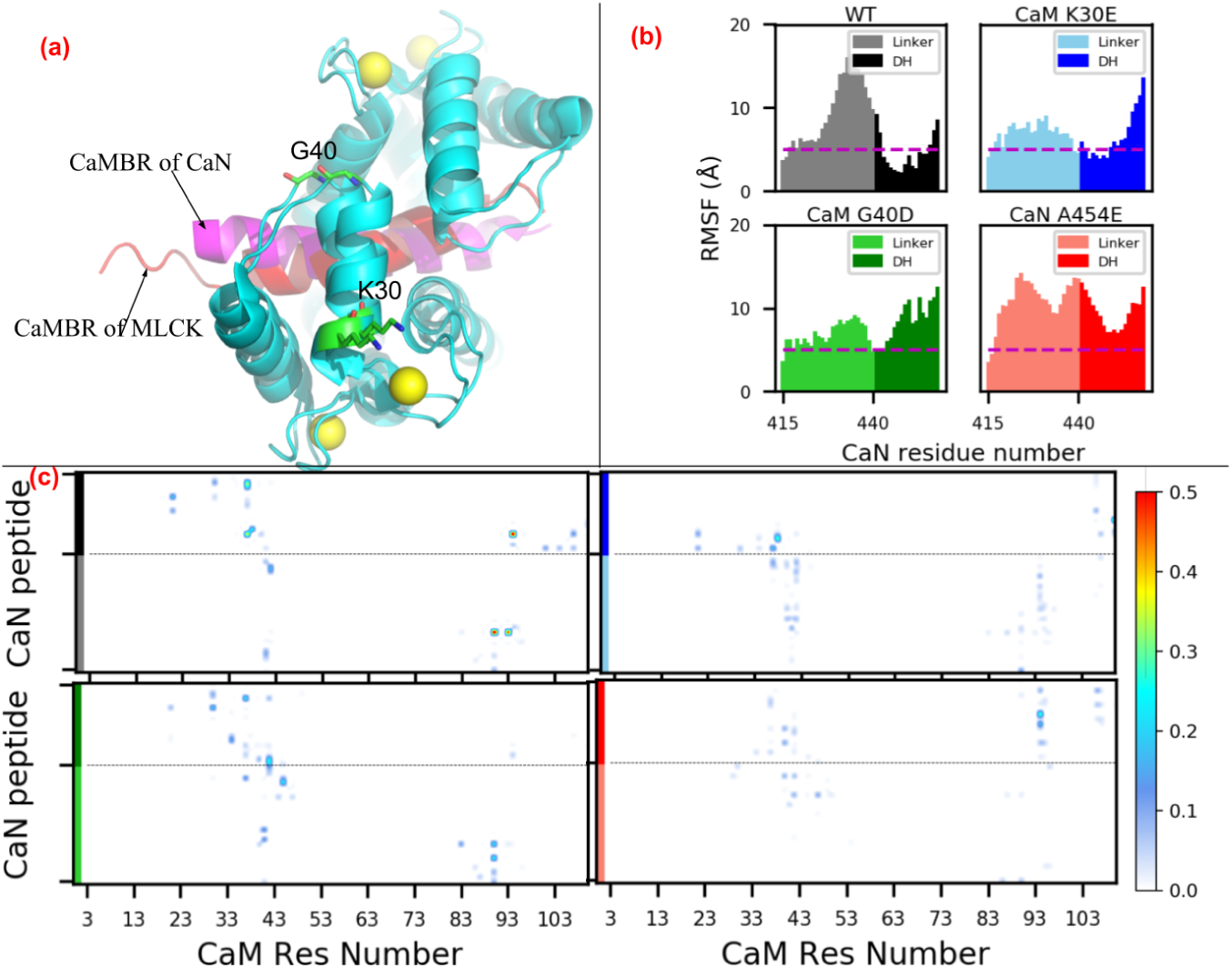
(a) Comparison of CaM-petide complex structure from CaN and MLCK (PDB ID: 2lv6 [67]). K30 and G40 are labeled (shown as sticks) based on their implication in the activation of the CaM target Myosin Light Chain Kinase (MLCK) [36] and proximity to site D determined by our simulations. (b) Non-hydrogen RMSF of linker and distal helix in WT and mutants. The dash line depicts RMSF value as 5 Å. (c) Percentage of simulated frames which have hydrogen bonds formed between linker/distal helix and CaM https://bitbucket.org/pkhlab/pkh-lab-analyses/src/default/2018-CaMDH/rmsf.ipynb and hbond.ipynb

We also reported the MM-GBSA-calculated binding free energies between distal helix and CaM of the mutants in Fig. 4. While the WT distal helix at the CaM site D has most stable binding with Δ*G* ≈ −27.7 kcal mol*^−^*^1^, the three mutations K30E, G40D and A454E have less favorable Δ*G*s as −21.8 kcal mol*^−^*^1^, −17.9 kcal mol*^−^*^1^ and −24.4 kcal mol*^−^*^1^ with P-values being 8.12 × 10*^−^*^2^, 5.1 × 10*^−^*^3^ and 2 × 10*^−^*^4^, respectively. The MM-GBSA-energies clearly shown that mutations would impair the binding affinity between distal helix and CaM. Accordingly, we presented linker and distal helix RMSF data for the WT and mutants in Fig. 9(b). The distal helix RMSF values among the two CaM variants were moderately increased compared to the WT case. Specifically, for the WT system, the distal helix residues were entirely within 10 Å and as low as ~ 2.5 Å. In contrast, the K30E variant yielded RMSF values no smaller than approximately 5 Å, while the C-terminal half approaches values nearing 15 Å. This trend manifested in fewer long-lived hydrogen bond contacts between the distal helix and both CaM domains (see Fig. 9). Similarly, the G40D mutation appeared to significantly disrupt interactions with CaN, as the entire distal helix region was characterized with RMSF values over ~10 Å in amplitude, with corresponding decreases in hydrogen bond contacts. We reported the MMGBSA calculated binding free energy between the distal helix and the CaM/CaMBR in Fig. 4. Among the mutations we considered, the A454E mutant had the most severe impact on RMSF values, as all residues comprising the linker and distal helix regions resulted in fluctuations above 8 Å. We also reported the *α*-helix probability of distal helix residue for variants in Fig. S7. It was found that all variants preserved a significant degree of overall helicity despite evidence of impaired interactions with CaM. However, the specific residues which formed *α*-helix were different among the variants: the mutation of A454 to E454 shifted the helicity to the first half of distal helix while the two CaM variants had the second half region being *α* helical. Altogether, these simulation data suggest that 1) the WT distal helix is stabilized at the site D CaM region, 2) site D residues R37 and K30 are implicated in distal helix binding and 3) disruption of site D binding by CaN A454E is consistent with reduced helicity and enzyme activity measured experimentally.

### 3.4 Phosphatase assays of site-directed CaM mutants

To support the simulation results, namely that the distal helix region binding predominantly to site D would impact CaN activity, we analyzed the kinetics of CaN mediated hydrolysis of pNPP. Our hypothesis was that disruption of site D/distal helix binding would reduce the accessibility of the catalytic site for pNPP binding which would reduce the apparent substrate affinity. This reduction would arise from the AID competing for the catalytic site, as a result of compromised site D/distal helix interactions. We therefore conducted pNPP assays using two site D variants, K30E and G40D. We analyzed substrate turnover in a Michaelis-Menten model, as described in the Methods. Phosphatase assays performed on CaM variants strongly suggest a statistically significant reduction (p-values in Table 1) in catalytic activity by a substantial increase in *K_M_* for K30E and G40D over the WT (27.6±1.1 mM, 46.0±2.3 mM, and 35.5±3.4 mM, respectively) indirectly indicating weaker binding of the distal helix peptide to the mutated CaM construct.

**Table 1:** Kinetic parameters of pNPP dephosphorylation with WT CaM and two site D variants. P-values given by Welch’s t-test for difference of means with unequal variance.

| CaM | $K_M$ (mM) | SD | p-value | $V_{max}$ ($\mu\text{mol}/\text{min}/\text{mg}$) | SD | p-value |
| --- | --- | --- | --- | --- | --- | --- |
| WT | 27.6 | 1.3 | - | 0.01 | 0.001 | - |
| K30E | 46.0 | 2.8 | 0.002 | 0.02 | 0.002 | 0.10 |
| G40D | 35.5 | 2.2 | 0.008 | 0.02 | 0.010 | 0.09 |

## 4 Discussion

### 4.1 Summary of Key Findings

We have used computational modeling to elucidate a potential mechanism for CaM-dependent regulation of CaN activity, through binding of a ‘distal helix’ region of the regulatory domain. Our microsecond-duration MD simulations indicate that the distal helix region maintains significant *α*-helical when bound to WT CaM. In contrast, we predict that an engineered variant (A454E) disrupts the domain’s secondary structure and ability to competently bind CaM. Both predictions are in agreement with experimental probes of CaN regulatory domain structure and phosphatase activity [1]. Namely, among the four potential regions on CaM’s surface that were solvent-accessible after binding the CaMBR, our data suggest that an RD region spanning the CaMBR through the distal helix was best stabilized at a site nestled between the CaM N- and C-terminal domains. In silico mutagenesis of two N-terminal CaM residues (K30E and G40D), prevented distal helix binding in our model, which we suggest hinders CaN activation, similar to identical mutations in CaM that were found to inactivate another CaM target, Myosin Light Chain Kinase (MLCK). We additionally report that our REMD simulations suggest that the isolated distal helix region spontaneously assumes significant *α*-helical in absence of CaM; which differ from trends observed in the complete RD domain observed experimentally [14]. We discuss this limitation and its implications in Sect. 4.5. Finally, we confirmed the potential CaM site D binding site for the distal helix through site-directed K30E and G40D variants, which we found to weaken CaN binding as reflected by an increase in *K_M_*(from 27.6 mM to 46.0 and 35.5 mM, respectively) in a pNPP phosphatase assay.

### 4.2 Plausible binding modes for putative CaN distal helix with CaM

Previous studies suggest that binding of regulatory domain residues beyond the CaMBR region are involved in CaM-dependent relief of CaN autoinhibition [1, 14]. Increases in regulatory *α*-helical content were reported upon binding CaM, that could not be accounted for by the CaMBR alone. Alanine to glutamic acid mutations at RD positions (A451E, A454E and A457E) C-terminal to the CaMBR decreased *α*-helical content and CaN activity. Further, HXMS studies indicate that in a complex of CaM with a regulatory domain/AID/C-terminal domain CaN construct that the CaMBR through distal helix regions had reduced solvent accessibility, suggestive of secondary interactions beyond the CaMBR. We calculated the backbone hydrogen bonds formed within the linker and distal helix region as indicator of solvent-protection and compared against experimental HXMS data. As shown in Fig. 10, In WT, site D has 16 hydrogen bonds, the 2 dominant hydrogen bonds (red arrow) are formed within the beta-sheet region (Fig. S5). Also in the distal helix region, two long-lived hydrogen bonds (>40% simulation time) exist. Compared with other cases, backbone hydrogens at site D would be most protected from HXMS due to the larger number of hydrogen bonds and relatively longer duration. Although A454E has largest number of hydrogen bonds, most are short-lived and the residue pairs which form hydrogen bonds are well seperated in sequence, indicating these hydrogen bonds do not contribute to *α*-helix secondary structure. Our computational modeling suggest that the putative distal helix region contains significant *α* helical character when bound to CaM site D, which qualitatively resemble those of experiment, suggesting reduced susceptibility to hydrogen/deuterium exchange. Nevertheless, compared to experimental HXMS data showing solvent-protected hydrogens are present across the whole linker and distal helix region, our computational backbone hydrogen bonds data indicates less degree of solvent-protection as the majority of hydrogen bonds are present in the N terminus of linker region and C-terminus of distal helix region in site D. This discrepancy could possibly be explained by the different lengths of CaN constructs used in HXMS experiment and our simulations. The construct in HXMS experiment contains the whole RD domain plus AID and C-tail while our simulations contain residues of A391 to I458 of RD domain. As shown in Fig. S9, we started from site D and run extra simulations with AID fragment added to distal helix region, the data clearly shows that the presence of extra residues would enhance the number and duration of backbone hydrogen bonds present in the linker and distal helix region.

**Figure 10:**
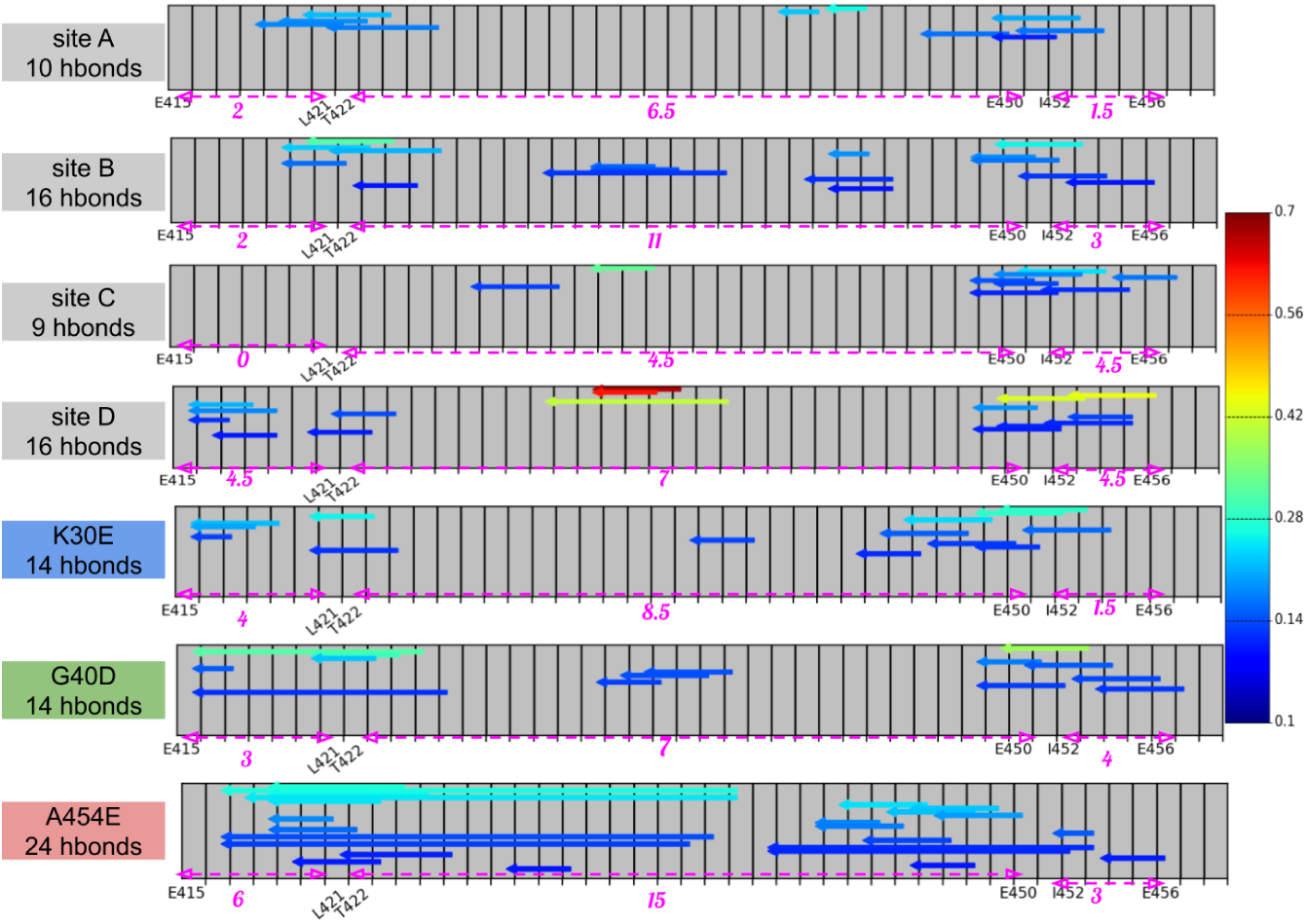
Backbone hydrogen bond analysis in the linker and distal helix region (E415 to I458). Each arrow represents one hydrogen bond with color indicating percentage of simulated frames with this hbond existed. Only hydrogen bonds exist >10% of simulation time are shown (we also show hdrogen bonds with >5% in Fig. S8). The whole region was divided into three subregions as indicated by the dashed magenta arrows below each subpanel. The subregion definition is consistent as the experimental HXMS data in Figure 8 in [14]. The number under the magenta arrow depicts the number of hydrogen bond in this subregion (one trans-subregion hydrogen bond contributes 0.5 to each subregion). https://bitbucket.org/pkhlab/pkh-lab-analyses/src/default/2018-CaMDH/analyze the backbone Hbond.ipynb

Additionally, several long-lived hydrogen bonds between the distal helix and CaM site D were found to stabilize the bound configuration, which dampened the variations of peptide position found at other identified sites (A-C) as reported by RMSF and energetic analyses (Fig. 8).

While we believe site D is the most probable site for distal helix binding, interactions with other potentially less-favorable sites could occur and contribute to the bound RD conformational ensemble. Such a diverse ensemble of strongly and weakly bound conformations is increasingly evident in complexes involving intrinsically disordered peptide (IDP)s and globular targets [27, 68] and may be adopted by CaN as well. It is also interesting that CD experiments in [1] suggested that the distal helix contact is abolished at temperatures above 38 degrees Celsius. It is tempting to speculate that the comparatively larger RMSFs of the bound distal helix configurations relative to the CaMBR, in addition to the weaker interaction energies, may render the distal helix interaction susceptible to melting.

Strengthening the case for the involvement of the CaM site D in binding the CaN distal helix are our comparisons against two CaM variants with substantially impaired ability to relieve enzyme auto-inhibition in another CaM target, Myosin Light Chain Kinase (MLCK) [36]. CaM appears to relieve MLCK auto-inhibition [69] through binding the kinase’s regulatory domain [70] and adopts a similar conformation as the CaN/CaM complex with CaM ‘wrapping’ around an *α* helical CaMBR motif (see also Fig. 9(a)) [1, 67]. Importantly, both appear to utilize secondary interactions beyond the CaMBR motif and it was shown by Van Lierop *et al* for MLCK that K30E and G40D mutations far from its CaMBR-binding domain prevented CaM-dependent kinase activity. These sites are localized to the site D region we identified for the distal helix in our study. Although the secondary interactions in MLCK likely involve CaM binding directly adjacent to the enzyme’s catalytic domain [71], we speculated that mutagenesis of these CaM residues could also impact CaN activation. Namely, we hypothesized that mutated these residues would destabilize distal helix binding. We confirmed this in our computational model by demonstrating less favorable distal helix binding scores.

### 4.3 Assessment of phosphatase activity

To challenge our hypothesis that impaired distal helix binding to CaM reduce CaN activity, we used kinetic phosphatase assays with the substrate pNPP on WT and the aforementioned CaM mutants. The Michaelis constant, *K_m_*, obtained from these experiments informs on the ability of the catalytic site to bind and dephosphorylate pNPP. This substrate is specific to the catalytic site due to its low molecular weight, which allows for an indirect analysis of the extent to which CaM binding removes the AID. We reported significantly higher *K_M_* for both K30E and G40D, thus these mutants evidence weaker distal helix binding that impedes removal of the AID from the CaN catalytic site. As a result, the CaM variants reduce the CaN catalysis of the dephosphorylation reaction, which can be interpreted as the AID competing within pNPP at the catalytic site and yielding a reduced apparent substrate affinity. This loss in affinity coincides with 40% increases in *K_M_* reported for CaN A454E relative to WT CaN [1], which were attributed to impaired distal helix formation. It should be noted that the small pNPP molecular is a preferable candidate for assessing distal helix binding, as opposed to common peptide-based dephosphorylation targets like RII [47]. Namely, the phospho-peptide binds to a site outside the active site (the LxVP site), therefore its binding, and hence Km, would be unaffected by mutations in the distal helix region. Rather, the effects would only be evident in the estimated Vmax values. pNPP, on the other hand, binds directly to the active site. Mutations in the distal helix region that disrupt its folding and allow the AID to bind to the active site would result in reduced pNPP binding (higher Km), but no change in Vmax. This explanation has been used by earlier authors studying the inhibitory properties of the AID as a peptide [47].

### 4.4 Tether-model of CaM-dependent CaN activation

We recognize that a shortcoming of our modeling approach is that it is limited to simulations of CaM complexes with fragments of the CaN regulatory domain, whereas distal helix binding’s effects on CaN activity are coupled to the entire regulatory domain and specifically, the AID. We therefore discuss a qualitative description of ‘linker’ dynamics of the regulatory domain appropriate for the AID-dependent inactivation of CaN. Specifically, we speculate that we can describe extents of CaN inactivation based on the AID’s effective concentration at the CaN catalytic site. This effective concentration is controlled by the tethering of the AID to CaN, which effectively confines the AID to a smaller volume (than free diffusion) that results in a higher interaction probability with the active site [72]. We use this effective concentration perspective to qualitatively assess how distal helix interactions with CaM impact CaN activity, as explicit all-atom simulations of the complete RD are prohibitively expensive. Here we leveraged previous theoretical models of protein activation [73, 74] by describing AID binding to the CaN catalytic domain as an intra-PPI. This PPI leverages a molecular tether (the regulatory domain) to enhances the *local* effective AID (*p*) concentration near the catalytic domain.

To illustrate this principle in CaN, we provide a basic extension of a linker-dependent modulation model we recently applied to the calcium-dependent troponin I (TnI) switch domain binding to troponin C (TnC) [72]. For this reaction, Ca^2+^ binding to TnC generates a conformation that can facilitate TnI binding:

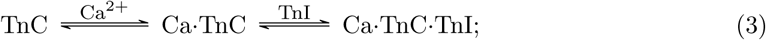

hence, increasing the TnI concentration would promote the generation of TnC·TnI with fewer equivalents of Ca^2+^. In the tethered state, we estimated that the *effective* switch peptide concentration was an order of magnitude greater near its TnC target than would be expected for a 1:1 stoichiometric ratio of untethered (free) switch peptide to TnC. Accordingly, we experimentally confirmed that formation of the TnC/TnI switch peptide occurred at lower Ca^2+^ concentrations for the TnC-tethered TnI compared to a cleaved system in which both TnC and TnI were untethered.

In a similar vein, we created a hypothetical linker-based model of CaN activation, based on a polymer-theory based model for the probability distribution of the linker spanning the CaMBR and AID domains (see Fig. 11). We introduce this model with several assumptions. Firstly, we postulate the CaN inhibition is dependent on the free AID concentration, of which the latter is determined by the RD ‘tether’ length. This tether length can assume three distributions associated with the CaM-free, CaMBR-bound CaM and CaMBR+distal helix-bound CaM, respectively. Lastly, for simplicity we assume that distal helix binds CaM independent of the AID’s bound state, though in reality we recognize there will be a competition between these two events.

**Figure 11:**
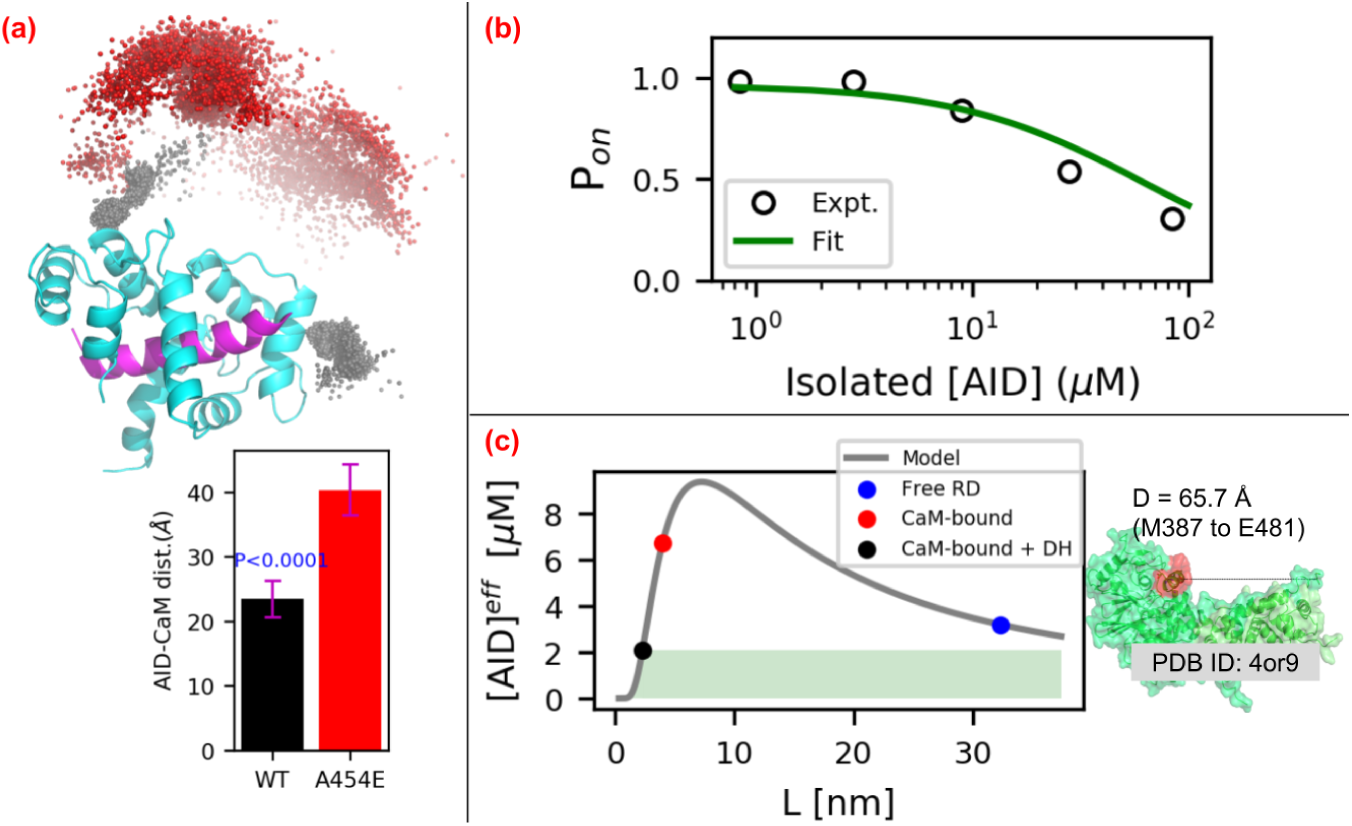
(a) Distribution of AID center of mass (COM) relative to the CaM/CaMBR complex. The black and red spheres represent the COMs of AID in WT and A454E cases, respectively. The lower panel depict distance between COMs of AID and CaM. The number above black bar are P values of WT case with null hypothesis that its values are equal to A454E case. (b) Fitting of the competitive-inhibitor model (Eq. 5) to experimental data from [75]. (c) Effective AID concentrations calculated via Eq. 4. The shaded green area represents effective [AID] that leads to CaN’s activation. Right panel illustrates the assumed distance between CaMBR and catalytic site. The value is set as 66 Å in this study. https://bitbucket.org/pkhlab/pkh-lab-analyses/src/default/2018-CaMDH/CaN_tether_model.ipynb

Under these assumptions, we describe the effective [AID] at the CaN catalytic domain, based on the RD linker length in its CaM-free, CaMBR-bound CaM and CaMBR+distal helix-bound CaM states. We based this on an effective concentration model for tethered ligands suggested by Van Valen *et al* [73],

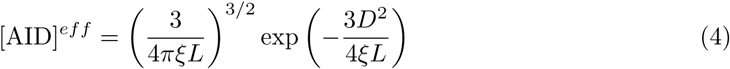

where *D* is the distance between CaMBR and catalytic site, *L* is linker length, and *ξ* is the persistence length. As stated in [73], although [AID]*^eff^* in Eq. 4 has unit of concentration, a real meaningful unit of [AID]*^eff^* that allows accurate interpretation of experiments could only be achieved via fitting to existing experimental data. Fortunately, experimental assays have been conducted to investigate the competitive inhibitory effect of isolated AID peptide on CaN phosphate activity on substrate peptide [75, 76]. In the assays, the reduction of phosphate activity was recorded as isolated AID peptide was added to intact CaN pre-incubated with CaM and substrate RII peptide. According to the experimental setup, there existed three competitive components that could bind the catalytic site of CaN: substrate RII peptide, isolated AID peptide and tethered AID from the intact CaN itself. Similar to *P_on_* definition which represents the probability of switch peptide being on under the competitive binding of free ligand and tethered ligand to receptor in [73], we also defined a *P_on_* which represents the percentage of CaN phosphate activity on substrate RII peptide under competitive binding from isolated AID peptide and tethered AID:

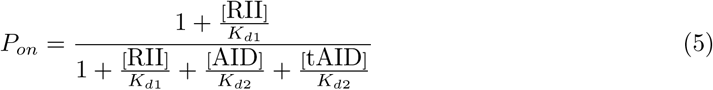

where [RII], [AID] and [tAID] are concentrations of substrate peptide, isolated and tethered AID peptide, respectively. [RII] is set as 5 *µ*M according to exprimental setup and the dissociation constant of substrate peptide *K_d_*_1_ is assumed to be 10 *µ*M. Tethered AID peptide is assumed to have same dissociation constant *K_d_*_2_ as isolated peptide with value being experimentally measured as 40 *µ*M [75, 76]. The fitting of Eq. 5 to experimental data in [75] with [tAID] as free pameter is shown in Fig. 11(b). [tAID] was fitted as 2.07 *µ*M and this value is corresponding to [AID]*^eff^* of ‘CaMBR+distal helix-bound CaM’ case in our tether model. In following tether model analysis, the [AID]*^eff^* from Eq. 4 were scaled by [tAID] to give meaningful unit of effective AID concentration.

To conduct tether model analysis, we first provide a rough estimate for the linker length through simulations of residues E415-M490 C-terminal to the CaMBR (see Fig. 11(a)). Starting from WT/A454E site D simulations, an optimized fragment (residues K459 to M490) containing AID built by tleap was fused to the C-termini of distal helix in the representative structure of first two most populated clusters. The complete structures were resolvated and simulated for ≈ 0.7*µs* as that described in Sect. S1.1. These simulations indicate that the WT AID to CaM distance is approximately 23 Å, versus approximately 40 Å for the A454E variant that precludes distal helix binding.

Based on these data, in Fig. 11(c) we demonstrate the effective AID concentration over a range of ligand lengths (L), predicted from Eq. 4 assuming *D* = 66 Å for the distance between CaM and the CaN AID binding site and *ξ* = 3 Å [77]. The black dot represents the CaMBR/distal helix (DH)-bound case, which has a tethered ligand length estimated from our simulation of approximately 23 Å or roughly 8 free amino acids. The blue dot represents free RD, which has ligand length of 95 residues (M387 to E481). The red dot represents the CaMBR-bound (no distal helix interaction as for the A454E case, in this case, the tethered ligand length estimated from our simulation as 40 Å). Based on these linker lengths, the corresponding effective [AID] concentrations for CaMBR-bound (A454E) states were 6.76 *µ*M versus 2.07 *µ*M for the CaMBR/distal helix-bound case. For the free RD case, the effective [AID] is 3.20 *µ*M. This approximate model qualitatively captures the experimental trends in activity data reported in the literature [1, 78], namely that maximal CaN activation requires CaM binding.

There are several considerations that could improve the accuracy of this model. These include assumptions that the linker follows a random-walk chain distribution, that the catalytic domain does not attract and thereby bias the AID distribution and that the CaN molecule does not sterically clash with the linker chain. Further, precise knowledge of the CaM distribution relative to the CaN B-chain would be needed to refine the effective linker lengths. Despite these assumptions, the model provides a qualitative basis for how RD mutations or variations in RD length could influence the efficiency of CaN (in)activation, similar to the model systems with synthetic linkers, as in [79].

### 4.5 Limitations

We observed appreciable degrees of alpha helical and beta sheet character in the regulatory domain that were not evident in the CD data from [14]. A primary distinction between the modeling and experimental studies is that we used a much smaller regulatory domain fragment (residue A391 to I458) than the full length domain in Rumi-Masante *et al* [14], owing to the computational expense. It is possible that there are different tendencies to form secondary structure, based on the regulatory domain length. Since we simulated only a small fragment of the RD domain, this might have increased the peptide’s preference for alpha helical structure than would otherwise be observed in measurements of the entire RD. For instance, it has been shown that IDPs have length-dependent preference of residue compositions as longer IDP has more enriched K, E and P than short IDP [80], implying the conformational properties of IDPs which are determined by sequence charge distribution [81] are also length-dependent. As a concrete example, Lin *et al* [82] reported that the 40-residue disordered amyloid beta monomer has reduced *β*-hairpin propensity when compared to the longer 42-residue monomer.

We additionally recognize that differences in ionic strength or solvent composition might influence the percentage of alpha helical character, although this seemed to be a modest effect in our simulations of the CaMBR alone [27]. Importantly, in that study, we reported negligible alpha helical character for that isolated CaMBR peptide, which suggests that our force field was not artificially stabilizing alpha helices, as had been an issue in earlier modeling studies of IDPs [83, 84]. Nevertheless, the potential overestimate of alpha helical content for the isolated peptide is probably of little consequence, since the predicted bound distal helix was shown to confirm exhibit significant alpha helical content consistent with experiment.

We utilized a REMD approach to sample the distal helix sequence in the absence of CaM; although REMD has been shown to perform well in terms of qualitatively describing conformational landscape, chemical shifts, *α*-helix stability for peptides of lengths comparable to the distal helix [85–87], we did not have the means to experimentally validate the predicted apo ensembles. Nevertheless, the simulations provide testable hypotheses in terms of the alpha helical content. We additionally limited ourselves to subsets of the CaM surface for the docking search, which represented approximately 38% of the solvent-exposed surface area. However, given that the microsecond-length simulations were sufficient to reorient the site B configurations into the site D site, we anticipate the docked distal helix candidates reasonably sampled the thermodynamically-accessible regions of the CaM surfaces. Although it has been demonstrated that RD binding to CaM is diffusion-limited, it is also possible that the intermediate complexes could be further optimized to form a final bound state, which would perhaps lead to more accurate assessments of critical intermolecular contacts and energy estimates. For the latter, alchemical methods such as thermodynamic integration may provide more accurate affinity estimates, albeit at a substantially greater computational expense compared to ‘end point’ methods like MM/GBSA. Lastly, more detailed simulations of the RD ensemble in the presence of the complete CaM and CaN structures are needed to more accurately characterize the effective AID distribution controlling CaN (in)activation.

## 5 Conclusions

We have developed a computational strategy to elucidate potential binding poses for a secondary interaction (the ‘distal helix’) between the CaN regulatory domain and CaM that is apparently essential for competent CaN activation. We combined REMD simulations of isolated distal helix peptides, protein-protein docking of the distal helix peptides to the CaMBR-bound CaM surface, and microsecond-scale MD simulations of candidate poses to implicate a so-called CaM site D in binding the CaN distal helix. The predicted site D region is in part stabilized through direct interactions with K30 and indirectly through G40, which is consistent with experimental probes of a CaM-activated enzyme, MLCK. With these data, we provide a qualitative model of AID-dependent CaN activation, which can be used to further refine potential molecular mechanisms governing the activation process and susceptibility to missense mutations. Given the broad range of physiological processes mediated by CaM binding to intrinsically disordered target proteins [60], the mechanistic details of CaN activation in this study may extend to diverse systems, including channel and cytoskeletal regulations [60, 88].

There are several compelling directions to pursue that would provide essential clues governing CaM-dependent CaN activation. For one, we have predicted several contacts that appear to be involved in stabilizing the distal helix region; mutagenesis of these potential ‘hotspots’ on the CaM and measurements of subsequent CaN phosphatase could help validate this site. In addition, more detailed characterization of the RD intrinsically-disordered conformation ensemble would benefit future modeling. Given the difficulty in probing ensemble properties of IDPs, it is likely that modeling and experiment, such as fluorescence resonance energy transfer (FRET) labeling, should work in tandem toward this goal. Furthermore, relating these RD ensemble properties to the propensity for AID and CaN catalytic domain interactions would comprise an essential step toward a complete model of CaM-dependent CaN activation.

## 6 Acknowledgement

We dedicate this study to the late Jeffry Madura, Ph.D., whose contributions to computational chemistry and the scientific community as a whole will be forever cherished. Research reported in this publication, release was supported by the Maximizing Investigators’ Research Award (MIRA) (R35) from the National Institute of General Medical Sciences (NIGMS) of the National Institutes of Health (NIH) under grant number R35GM124977. This work used the Extreme Science and Engineering Discovery Environment (XSEDE) [89], which is supported by National Science Foundation under grant number ACI-1548562.

## S1.1 Methods

The initial structure corresponding to CaN distal helix to AID region (residues 459 to 490) was built from sequence via tleap. The initial structure was subjected to minimization and MD simulation in vacuum according to the procedure described Sect. 2.3. The optimized structure was then appended to the C-terminus of distal helix region from the representative structure of site D simulations via tleap. The representative structures of the first two most populated clusters from site D simulations were selected, making the simulation duplicate. The sytem was then solvated in TIP3P waterbox with 0.15 M KCl ions added. The simulation details are same as previous section in which the tleap built structure was first relaxed while rest part being fixed during the heating and equilibrium stage. After reaching equilibrium, about 0.7 *µ*s production simulation was performed from each replica of the duplicate. The simulations were repeated for the CaN A454E mutant. The total number of MD cases considered in this study is listed in Table S3.

## S1.2 Tables

**Table S1:** Residues at each tentative binding site on collapsed CaM used in ZDOCK to predict distal helix interaction at each site

| Tentative Site | Residues |
| --- | --- |
| Site A | R86, F89, V142, Y138 |
| Site B | R106, I125, D118, D122 |
| Site C | I9, F12, L69, F65 |
| Site D | K30, T34, R37, Q49 |

**Table S2:**
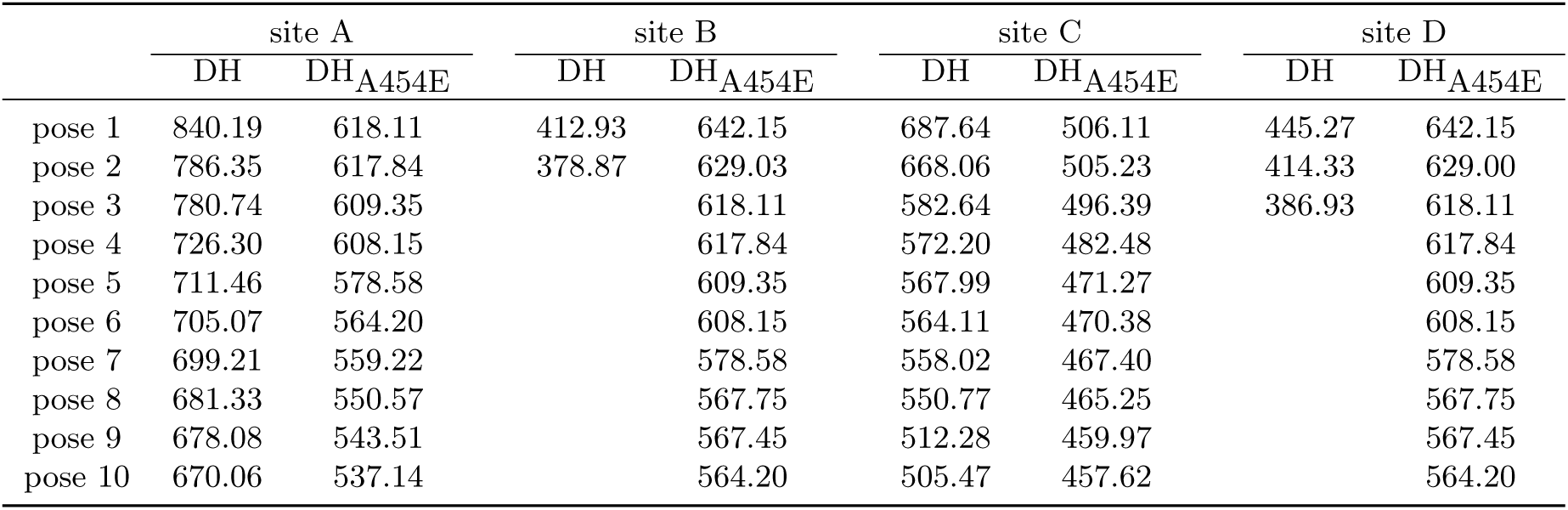
ZDOCK docking scores of distal helix and DH_A454E_ at each site. During each docking, a culling process was applied on the initially generated 2 × 10^3^ poses to eliminate those having no contacts with residues we specified in Table S1. After culling, the docking scores of highest-scored poses (up to 10 poses) are given. In DH-site A and DH_A454E_-site B cases, there was no remaining poses left after culling, we instead output the first 10 poses’ scores from the initially-generated 2 *×* 10^3^ poses.

|  | site A |  | site B |  | site C |  | site D |  |
| --- | --- | --- | --- | --- | --- | --- | --- | --- |
|  | DH | DH<sub>A454E</sub> | DH | DH<sub>A454E</sub> | DH | DH<sub>A454E</sub> | DH | DH<sub>A454E</sub> |
| pose 1 | 840.19 | 618.11 | 412.93 | 642.15 | 687.64 | 506.11 | 445.27 | 642.15 |
| pose 2 | 786.35 | 617.84 | 378.87 | 629.03 | 668.06 | 505.23 | 414.33 | 629.00 |
| pose 3 | 780.74 | 609.35 |  | 618.11 | 582.64 | 496.39 | 386.93 | 618.11 |
| pose 4 | 726.30 | 608.15 |  | 617.84 | 572.20 | 482.48 |  | 617.84 |
| pose 5 | 711.46 | 578.58 |  | 609.35 | 567.99 | 471.27 |  | 609.35 |
| pose 6 | 705.07 | 564.20 |  | 608.15 | 564.11 | 470.38 |  | 608.15 |
| pose 7 | 699.21 | 559.22 |  | 578.58 | 558.02 | 467.40 |  | 578.58 |
| pose 8 | 681.33 | 550.57 |  | 567.75 | 550.77 | 465.25 |  | 567.75 |
| pose 9 | 678.08 | 543.51 |  | 567.45 | 512.28 | 459.97 |  | 567.45 |
| pose 10 | 670.06 | 537.14 |  | 564.20 | 505.47 | 457.62 |  | 564.20 |

**Table S3:**
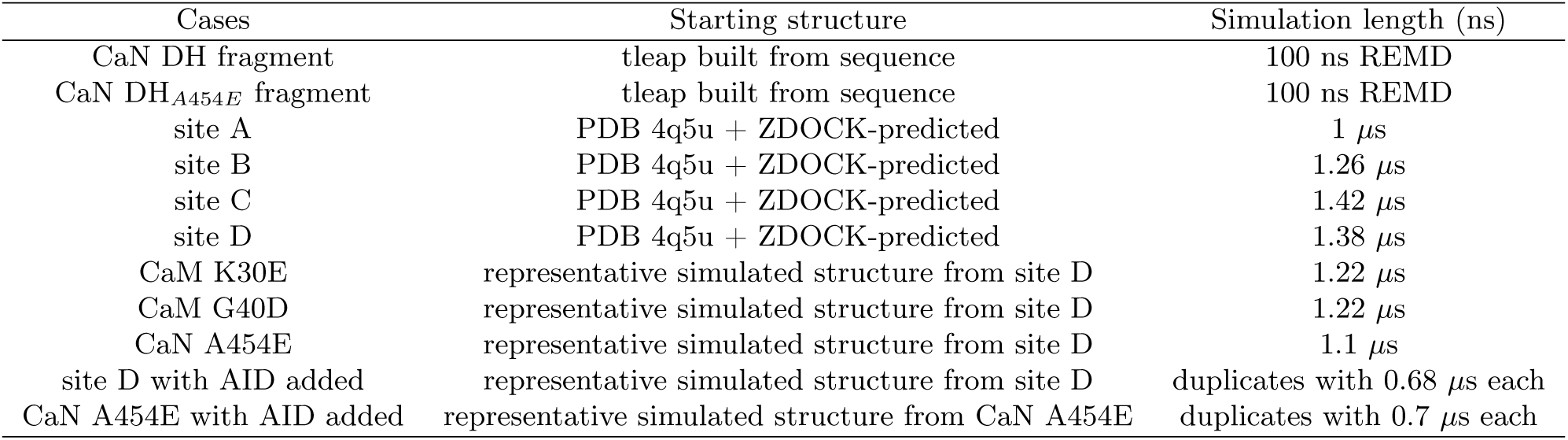
MD simulation cases.

| Cases | Starting structure | Simulation length (ns) |
| --- | --- | --- |
| CaN DH fragment | tleap built from sequence | 100 ns REMD |
| CaN DH<sub>A454E</sub> fragment | tleap built from sequence | 100 ns REMD |
| site A | PDB 4q5u + ZDOCK-predicted | 1 μs |
| site B | PDB 4q5u + ZDOCK-predicted | 1.26 μs |
| site C | PDB 4q5u + ZDOCK-predicted | 1.42 μs |
| site D | PDB 4q5u + ZDOCK-predicted | 1.38 μs |
| CaM K30E | representative simulated structure from site D | 1.22 μs |
| CaM G40D | representative simulated structure from site D | 1.22 μs |
| CaN A454E | representative simulated structure from site D | 1.1 μs |
| site D with AID added | representative simulated structure from site D | duplicates with 0.68 μs each |
| CaN A454E with AID added | representative simulated structure from CaN A454E | duplicates with 0.7 μs each |

**Table S4:** Hydrogen bonds between distal helix and CaM in each case. Only hydrogen bonds sustained for *>*= 10% of the simulation duration are listed. The first residue is from distal helix and second residue is from CaM

| Site A | Site B | Site C | Site D | CaM K30E | CaM G40D | CaN A454E |
| --- | --- | --- | --- | --- | --- | --- |
| Q442-R90 0.17 | K441-G40 0.10 | Q442 R90 0.17 | Q442 S101 0.10 | Q445 S38 0.10 | S446 T34 0.14 | V459 K94 0.18 |
| K441-R90 0.27 | A457 R37 0.10 | E453 L4 0.13 | E450 K21 0.10 | Q445 K21 0.10 | S446 N111 0.16 | A451 K94 0.28 |
|  | E445 N42 0.11 | E453 Q3 0.28 | E456 K30 0.11 | A454 K115 0.10 | E453 K30 0.19 |  |
|  | L444 N42 0.12 |  | Q445 H107 0.12 | D455 K115 0.15 | K441 N42 0.21 |  |
|  | E456 K21 0.13 |  | E453 K21 0.19 | Q442 R37 0.19 | D455 R37 0.22 |  |
|  | E456 S38 0.17 |  | S446 S38 0.23 | L444 S38 0.29 | Q442 N42 0.24 |  |
|  | K441 R90 0.23 |  | D455 R37 0.23 | T448 T110 9.35 |  |  |
|  | K441 D93 0.28 |  | E456 R37 0.32 |  |  |  |
|  | D455 R37 0.30 |  | Q445 R37 0.37 |  |  |  |
|  |  |  | Q445 K94 0.55 |  |  |  |

## S1.3 Figures

**Figure S1:**
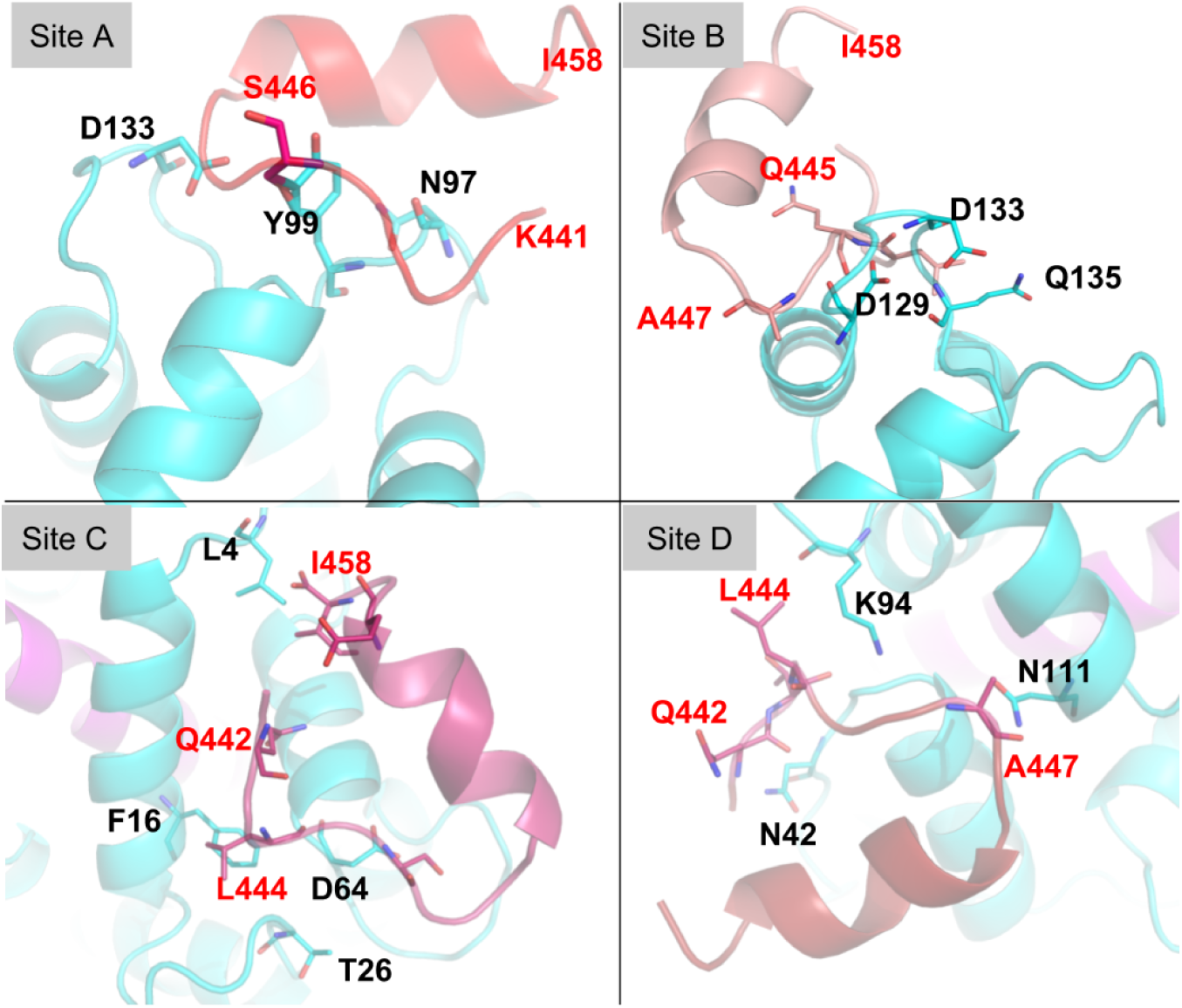
Highest-ranking CaM/distal helix interaction poses predicted by ZDOCK3.0.2 [31] webserver at each site. The color scheme is same as Fig. 2. Key residues at the interaction surface are shown in sticks with black labels for CaM residues and red labels for distal helix residues. Comparisons of the WT distal helix poses versus predictions for DH_A454E_ are shown in Fig. S2.

**Figure S2:**
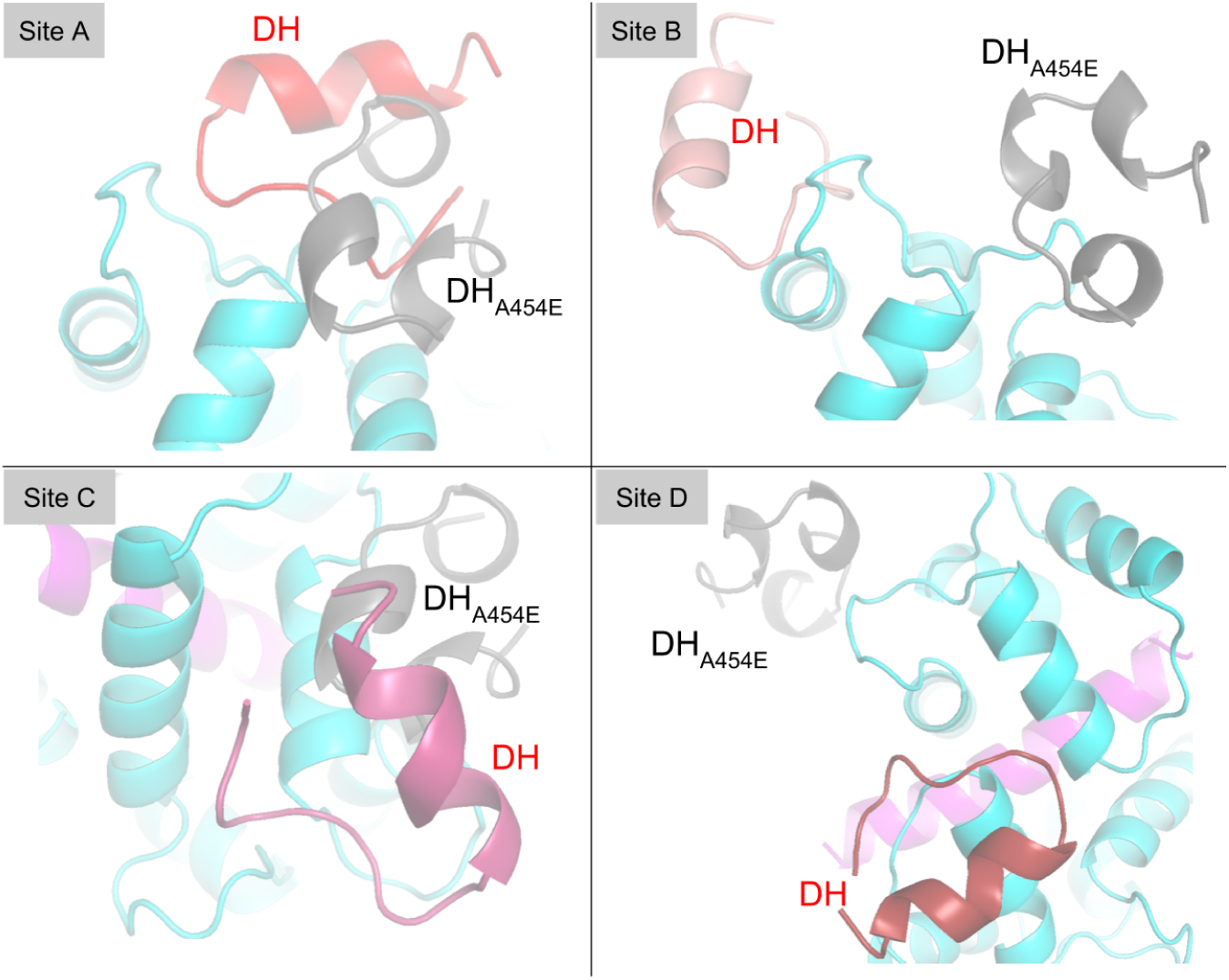
Comparison of Zdock predicted poses of DH and DH_A454E_ mutant at each site. The DH_A454E_ mutant is colored in gray. At site A and C, the poses of DH and mutant are close, while at site B and D, the DH_A454E_ mutant are predicted to be located near site A.

**Figure S3:**
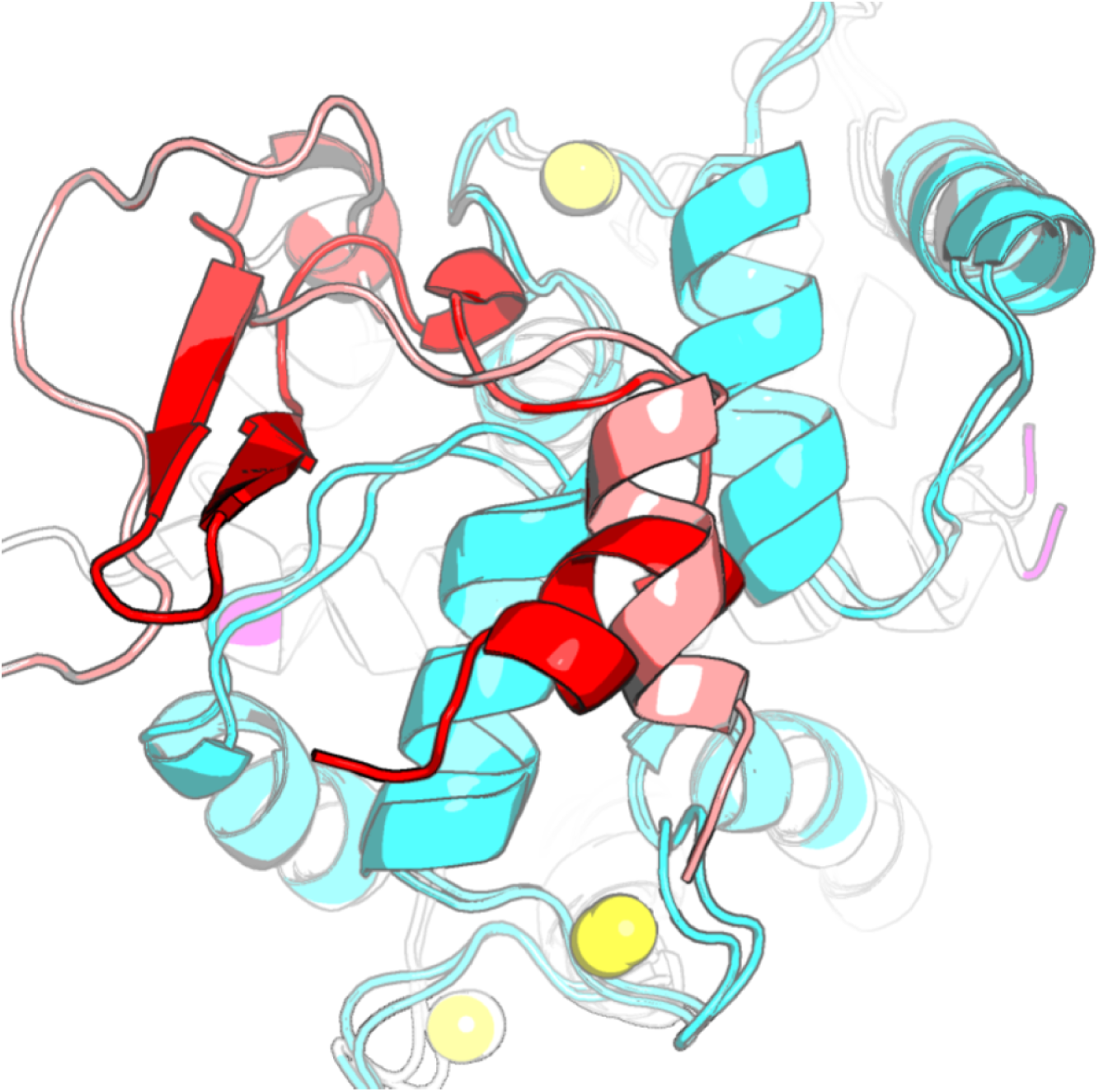
Overlap of MD simulated distal helix conformation starting at site B (colored in salmon) and site D (colored in red). During the simulations, distal helix starting at site B migrated to site D. go to /net/share/bsu233/CaMDH/zdock_MD/zdocksite4/traj/pdb/s4*−*s2*−*comparasion, type “pymol log.pml”

**Figure S4:**
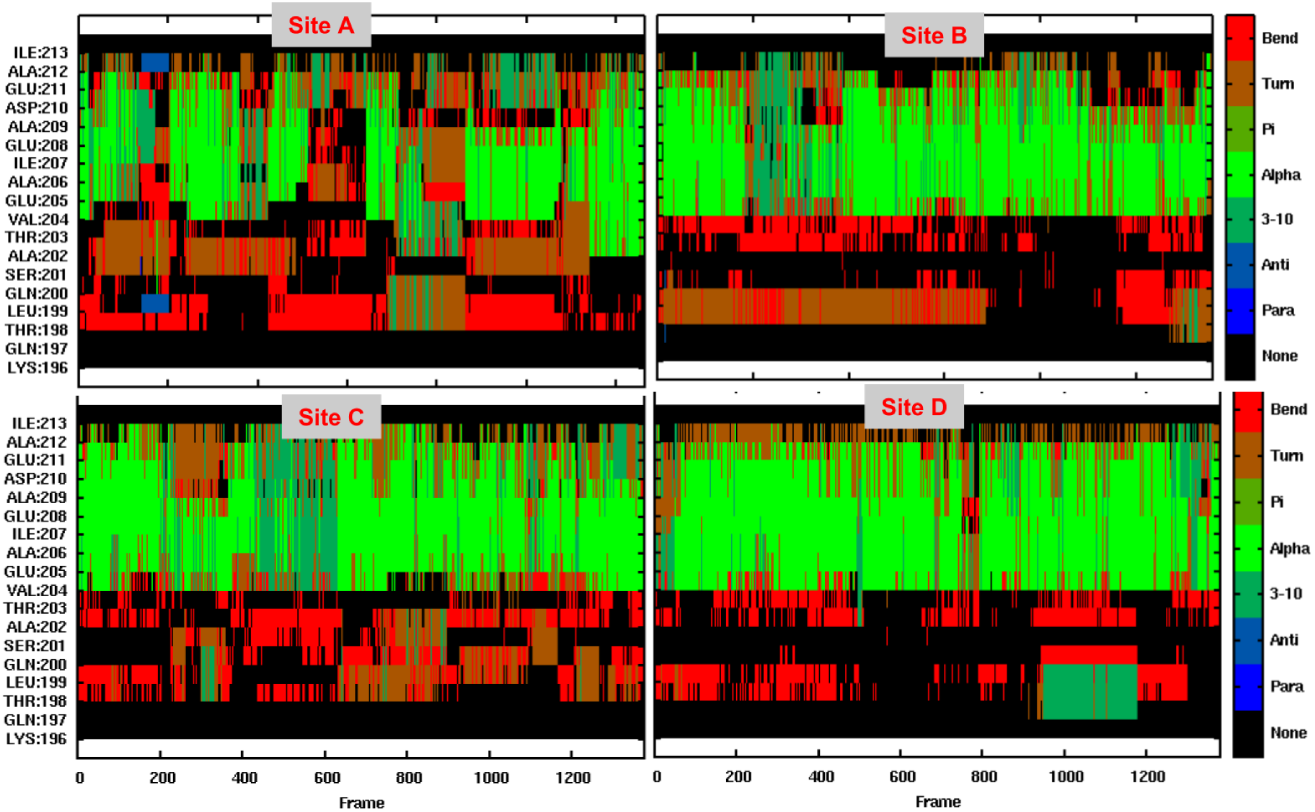
Secondary structure propensity of the distal helix during the MD simulations, as predicted via CPPTRAJ using the DSSP algorithm. The abundance of green indicates significant alpha helicity.

**Figure S5:**
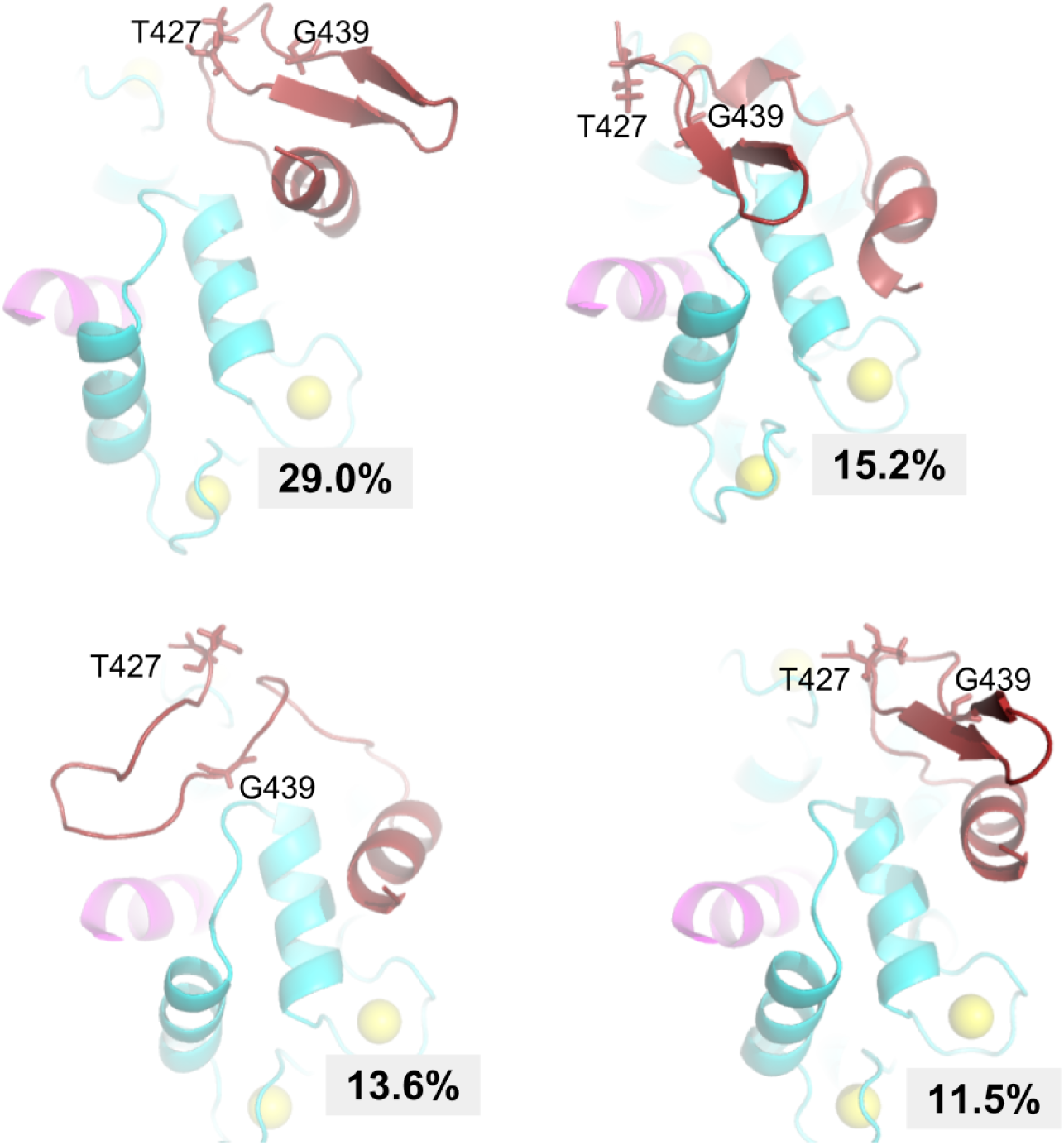
Illustration of *β*-sheet formed in T427-G439 region from site D simulations. The shown structures are representative structures of first four most populated clusters.

**Figure S6:**
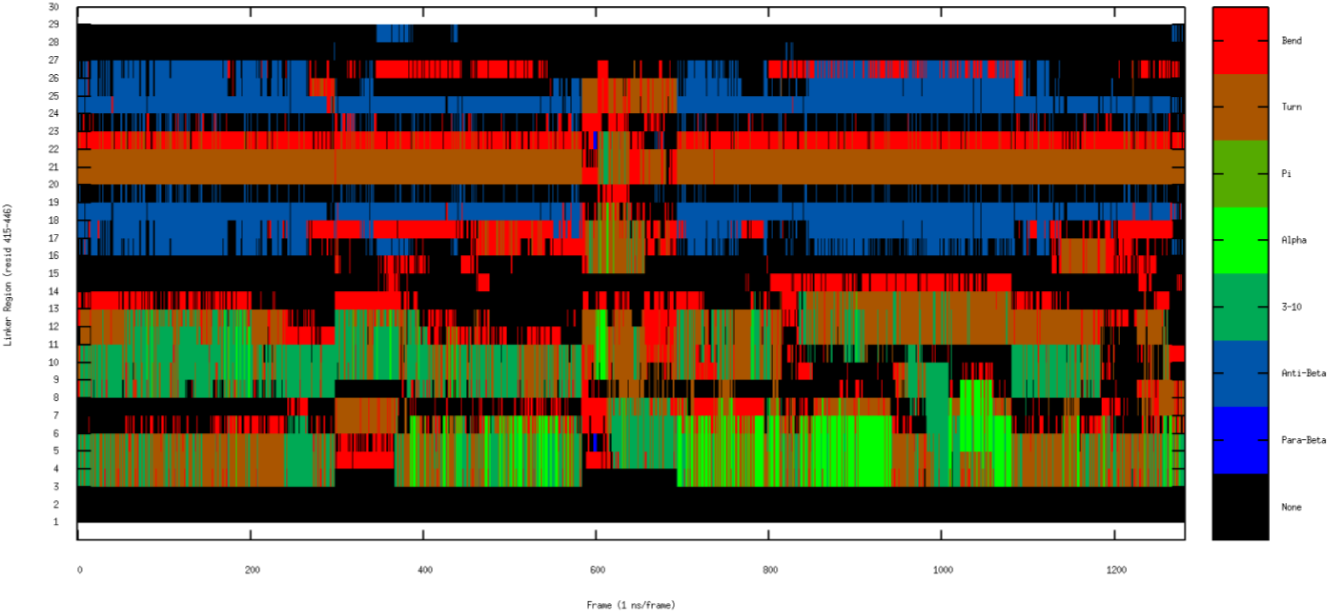
Secondary structure propensity of the linker at site D during the MD simulations, as predicted via CPPTRAJ using the DSSP algorithm. The blue region depicts the formation of *β*-sheet during most of the simulation time.

**Figure S7:**
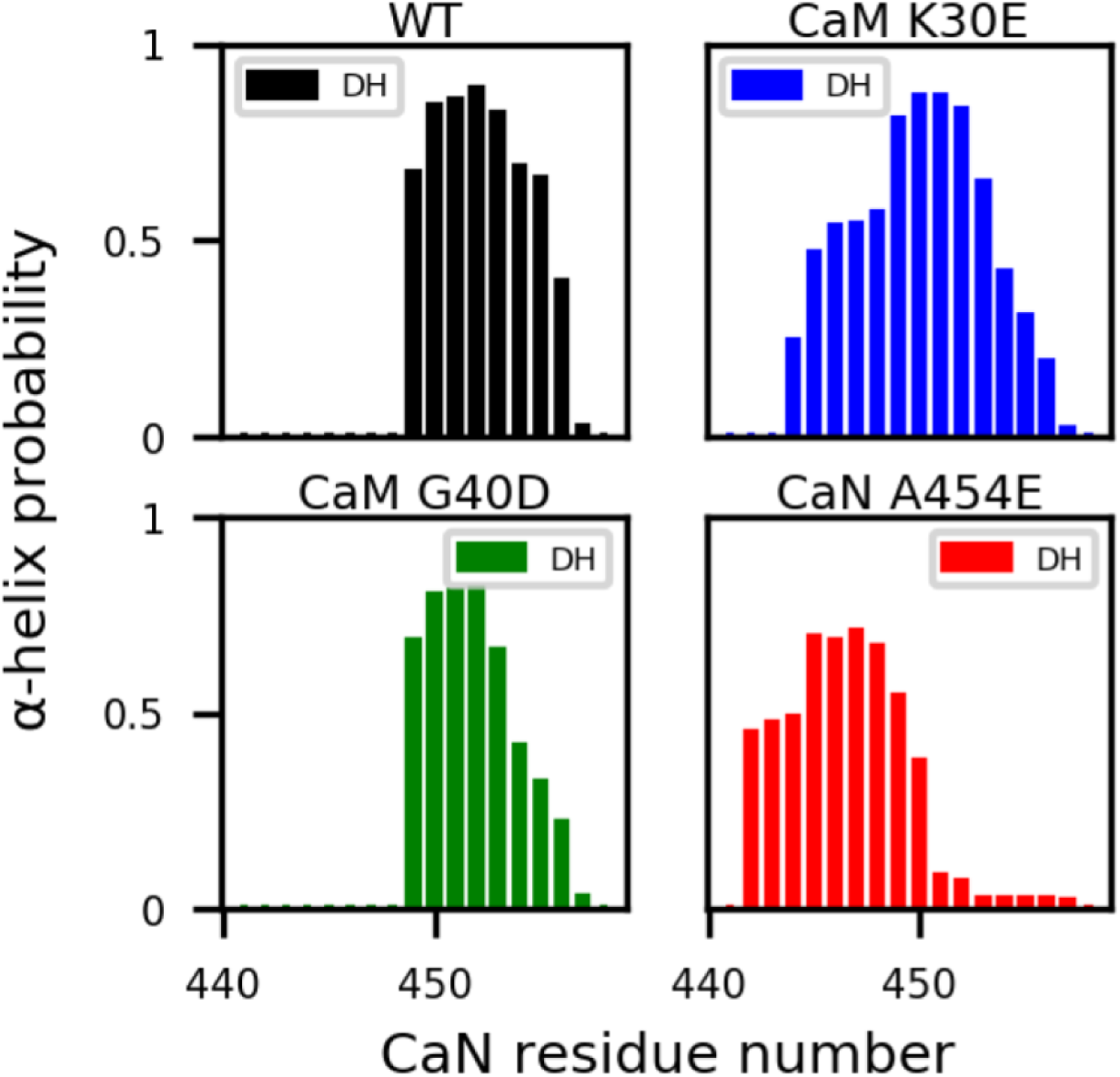
*α*-helix structural probability of each residue in distal helix region of WT, CaM K30E and G40D mutants and CaN A454E mutant calculated from MD simulations initiated at site D.

**Figure S8:**
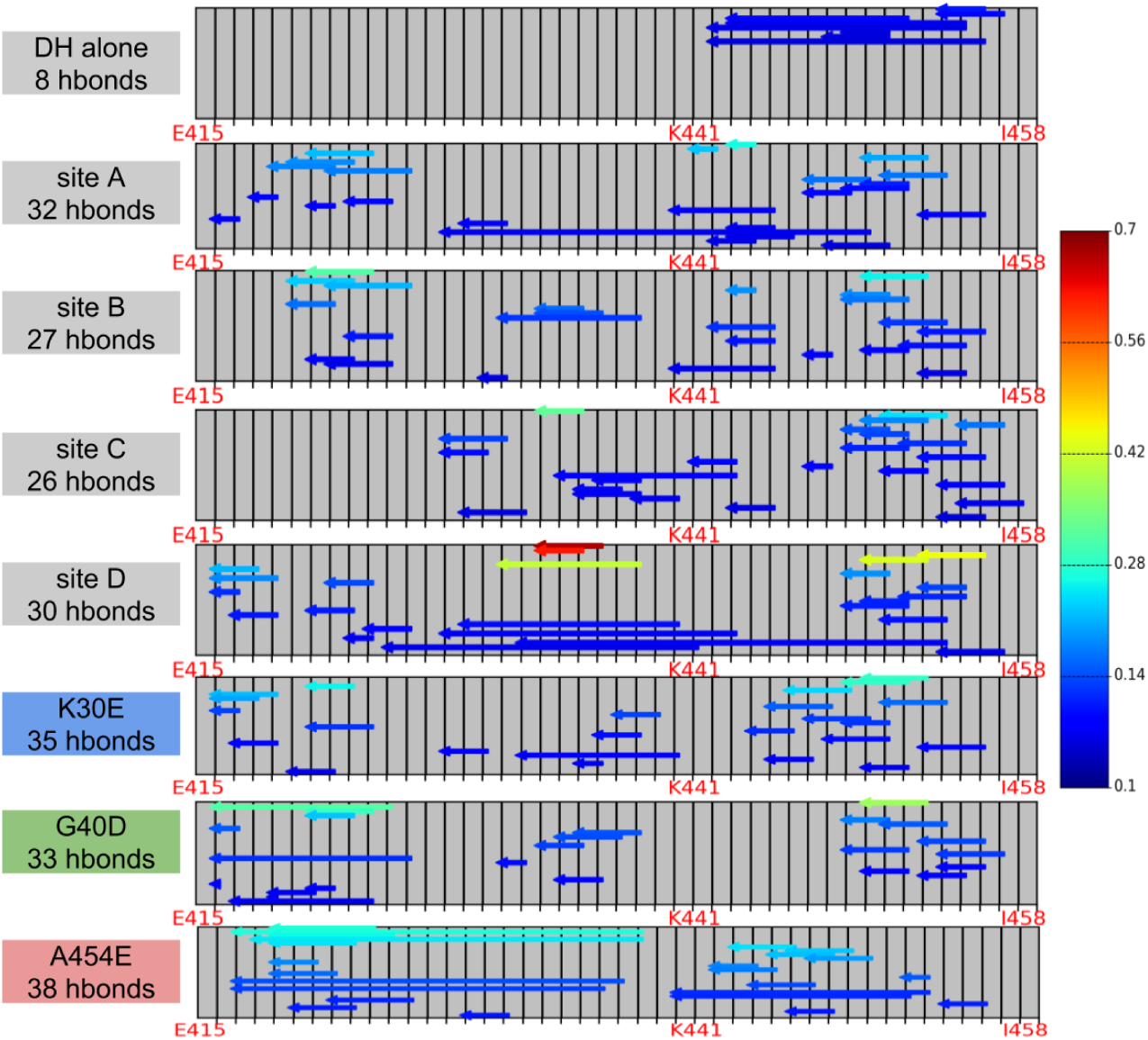
Backbone hydrogen bond analysis in the linker and distal helix region. Each arrow represents one Hbond with color indicating percentage of simulated frames with this hbond existed. hbonds exist >5% of simulation time are shown. https://bitbucket.org/pkhlab/pkh-lab-analyses/src/default/2018-CaMDH/analyze the backbone Hbond.ipynb

**Figure S9:**
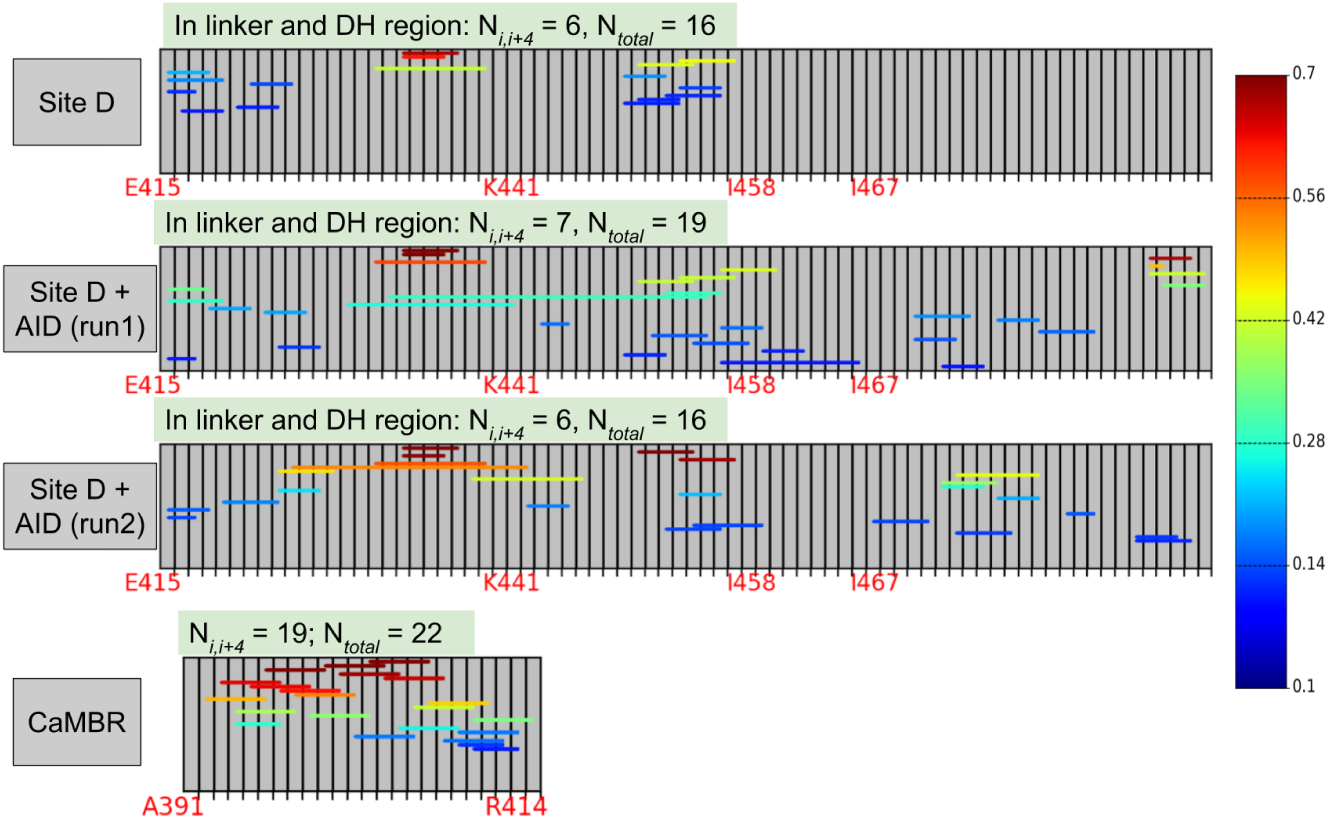
Comparison of backbone hydrogen bond in the linker and DH region between WT site D and WT site D with extra AID fragment added. The backbone hydrogen bond of CaMBR from WT site D is also show as reference. 1) In one of the two runs with AID added, the number of i,i+4 hbond (Ni,i+4) was increased by 1 due the presence of AID fragment, while the other run has value unchanged. 2) In both runs with AID added, the backbone hbonds become more stable as the duration of hbonds are generally enhanced. 3) The helicity of linker and DH region in site D (with/without AID) are much less stable than that of CaMBR, as indicated by smaller value of Ni,i+4, 6/7 vs. 19 https://bitbucket.org/pkhlab/pkh-lab-analyses/src/default/2018-CaMDH/analyze the backbone Hbond.ipynb

